# Alternative splicing of auxiliary β2-subunits stabilizes Cav2.3 Ca^2+^ channel activity in continuously active midbrain dopamine neurons

**DOI:** 10.1101/2021.02.10.430224

**Authors:** Anita Siller, Nadja T. Hofer, Giulia Tomagra, Nicole Wiederspohn, Simon Hess, Julia Benkert, Aisylu Gaifullina, Desiree Spaich, Johanna Duda, Christina Pötschke, Kristina Vilusic, Eva Maria Fritz, Toni Schneider, Peter Kloppenburg, Birgit Liss, Valentina Carabelli, Emilio Carbone, Nadine J. Ortner, Jörg Striessnig

**Affiliations:** Department of Pharmacology and Toxicology, Institute of Pharmacy, Center for Molecular Biosciences Innsbruck, University of Innsbruck, Innsbruck, Austria; Institute of Neurophysiology, University of Cologne, Cologne, Germany; Department of Drug Science, NIS Centre, University of Torino, Torino, Italy; Institute of Applied Physiology, University of Ulm, Ulm, Germany; Institute for Zoology, Biocenter, CECAD, University of Cologne, Cologne, Germany

**Author notes:** Correspondence: Nadine J. Ortner or Jörg Striessnig.

**Keywords:** Cav2.3, R-type Ca^2+^ channel, β-subunits, alternative splicing, dopaminergic neurons, Parkinson’s disease

## Abstract

In dopaminergic (DA) substantia nigra (SN) neurons Cav2.3 R-type Ca^2+^-currents contribute to somatodendritic Ca^2+^-oscillations. These may contribute to the selective degeneration of these neurons in Parkinson’s disease (PD) since Cav2.3-knockout is neuroprotective in a PD mouse model. However, the typical Cav2.3 gating would predict complete channel inactivation during SN DA neuronal firing. Here we show that in tsA-201-cells the membrane-anchored β2-splice variants β2a and β2e stabilize Cav2.3 gating properties allowing sustained Cav2.3 availability during simulated pacemaking and enhanced Ca^2+^-currents during bursts. We confirmed the expression of β2a and β2e-subunits in the SN and identified SN DA neurons. Patch-clamp recordings of SN DA neurons in mouse brain slices revealed R-type Ca^2+^-currents similar to β2a- or β2e-stabilized Cav2.3-currents and recordings in cultured murine DA neurons confirmed their activity during pacemaking. Taken together, our data support an important (patho)physiological role of β-subunit alternative splicing for Cav2.3 Ca^2+^-signaling in highly vulnerable SN DA neurons.

## Introduction

Parkinson’s disease (PD) is one of the most common neurodegenerative disorders. Its motor symptoms are characterized by progressive degeneration of dopamine (DA)-releasing neurons in the substantia nigra (SN), while neighboring DA neurons in the ventral tegmental area (VTA) remain largely unaffected (Damier et al., 1999; Giguère et al., 2018; Surmeier et al., 2017). Current PD therapy is only symptomatic and primarily based on the substitution of striatal DA by administration of L-DOPA or dopamine D2 receptor agonists. Unfortunately, none of the existing therapeutic approaches for PD patients is disease-modifying and can prevent disease progression (for review see Liss & Striessnig, 2019; Surmeier et al., 2011; Surmeier et al., 2017).

The development of novel neuroprotective strategies for the treatment of early PD requires the understanding of the cellular mechanisms responsible for the high vulnerability of SN DA neurons. Among these mechanisms elevated metabolic stress appears to play a central role (for review see Liss & Striessnig, 2019), eventually triggering lysosomal, proteasomal and mitochondrial dysfunction (Burbulla et al., 2017; Surmeier et al., 2017). Intrinsic physiological properties of SN DA neurons, in particular increased cytosolic DA levels and high energy demand due to large axonal arborisation favour metabolic stress (Bolam & Pissadaki, 2012; Liss & Striessnig, 2019). In addition, these neurons must handle a constant intracellular Ca^2+^-load resulting from dendritic and somatic Ca^2+^-oscillations triggered during their continuous electrical activity (Ortner et al., 2017; Surmeier et al., 2011). Dendritic Ca^2+^-transients largely depend on the activity of voltage-gated Ca^2+^ channels, in particular Cav1.3 L-type (LTCCs) and T-type channels (Guzman et al., 2018). Cav1.3 channels can activate at subthreshold membrane potentials (Koschak et al., 2001; Lieb et al., 2014; Xu & Lipscombe, 2001) and do not completely inactivate during continuous pacemaking activity (Guzman et al., 2018; Guzman et al., 2009; Ortner et al., 2017). Some, but not all, in vivo studies (Liss & Striessnig, 2019) showed neuroprotection by the systemic administration of dihydropyridine (DHP) L-type channel blockers in 6-OHDA and MPTP animal models of PD thus further supporting a role of LTCCs as potential neuroprotective drug target. Based on these preclinical data and supporting observational clinical evidence (Liss & Striessnig, 2019), the neuroprotective potential of the DHP isradipine (ISR) was tested in a double-blind, placebo-controlled, parallel-group phase 3 clinical trial (“STEADY-PD III”, NCT02168842; Biglan et al., 2017). This trial reported no evidence for neuroprotection by ISR. Several explanations have been offered for this negative outcome (Parkinson Study Group, 2020). One likely explanation is that voltage-gated Ca^2+^-channels (Cavs) other than LTCCs also contribute to Ca^2+^-transients in SN DA neurons. This is supported by the observation that only about 50% of the Ca^2+^-transients are blocked by isradipine in the dendrites of SN DA neurons (Guzman et al., 2018) and that action potential-associated Ca^2+^-transients in the soma appear to be even resistant to isradipine (Ortner et al., 2017). Therefore, in addition to L-type, other types of Cavs expressed in SN DA neurons (Branch et al., 2014; Evans et al., 2017; Philippart et al., 2016) may also contribute to Ca^2+^-induced metabolic stress in SN DA neurons. Cav2.3 (R-type) Ca^2+^-channels are very promising candidates. We have recently shown that SN DA neurons in mice lacking Cav2.3 channels were fully protected from neurodegeneration in the chronic MPTP-model of PD. Moreover, we found that Cav2.3 is the most abundant Cav expressed in SN DA neurons, and substantially contributes to activity-related somatic Ca^2+^-oscillations (Benkert et al., 2019). These findings make Cav2.3 R-type Ca^2+^ channels a promising target for neuroprotection in PD.

SN DA neurons are spontaneously active, pacemaking neurons, either firing in a low-frequency single-spike mode or transiently in a high-frequency burst mode (Grace & Bunney, 1984a, 1984b; Paladini & Roeper, 2014). During regular pacemaking their membrane potential is, on average, rather depolarized ranging from about −70 mV after an action potential (AP) to about −40 mV at firing threshold (Gantz et al., 2018; Guzman et al., 2018; Ortner et al., 2017). The contribution of a particular Cav channel to Ca^2+^-entry is therefore largely determined by its steady-state inactivation properties, which determines its availability at these depolarized voltages. In Cav2.3 channels expressed with auxiliary α2δ and various β-subunits steady-state inactivation occurs at much more negative potentials being almost complete at voltages positive to −50 mV. Moreover, Cav2.3 channels inactivate quickly during depolarizations (Jones et al., 1998; Pereverzev et al., 2002; Soong et al., 1993; Williams et al., 1994; Yasuda et al., 2004). Together these properties predict that most of the Cav2.3 channel current is inactivated during continuous SN DA neuron pacemaking.

Here we directly addressed this question by stimulating Cav2.3 channel complexes using command voltages simulating either SN DA neuron-like tonic pacemaking or brief burst activity. As previously described for LTCCs (Ortner et al., 2017) we expressed Cav2.3 channels together with auxiliary subunits in tsA-201 cells because individual Ca^2+^-current components are difficult to resolve in patch-clamp recordings of SN DA neurons in slices. We found that Cav2.3 currents rapidly decay when co-expressed with β3 and β4 subunits. However, Cav2.3 remained active during continuous pacemaking in the presence of the β2a and β2e splice variants, which stabilize steady-state inactivation of Cav2.3 at more positive potentials (up to 35 mV more positive compared to β3) and considerably slow current inactivation. In contrast, steady-state inactivation of Cav1.3 channels was largely unaffected by β2a, suggesting that Cav1.3 availability is much less dependent on the presence of β2a subunits. Using RNAScope we confirmed the presence of both, β2a and β2e transcripts in identified mouse SN and VTA DA neurons and quantitative PCR analysis showed that β2-subunits represent about 25% of all β-subunit transcripts in the SN and VTA with about 50% comprising β2a and β2e variants. In patch-clamp recordings of SN DA neurons in mouse brain slices we detected R-type Ca^2+^-currents similar to β2a- or β2e-stabilized Cav2.3-currents and recordings in cultured DA neurons confirmed R-type current activity during the pacemaking cycle. Taken together, our data further support a role of Cav2.3 in SN DA neuron Ca^2+^-signaling and reveal an important (patho)physiological role of β-subunit alternative splicing.

## Results

### SNX-482 inhibits Ca^2+^ current (I_Ca_) and reduces spontaneous AP firing in cultured mouse midbrain DA neurons

We have recently shown that in mouse SN DA neurons Cav2.3 channels contribute to action potential (AP)-induced somatic Ca^2+^-oscillations. In Cav2.3-deficient mice the amplitude of the Ca^2+^-oscillations was decreased by about 50%. This continuous Ca^2+^ load can potentially contribute to the high vulnerability of these neurons in PD (Benkert et al., 2019).

To further confirm the presence of functional Cav2.3 channels we investigated effects of the Cav2.3 blocker SNX-482 on firing frequency and AP shape in spontaneously firing cultured mouse DA midbrain neurons. We employed SNX-482 at a low concentration (100 nM) to inhibit Cav2.3 current components (IC_50_=30 nM) but spare L-type channels (IC_50_>1 µM, Bourinet et al., 2001; Newcomb et al., 1998). SNX-482 had profound effects on firing properties. In current-clamp recordings it significantly reduced the spontaneous firing frequency from 4.1 ± 0.8 Hz (control, n=10, N=3; 95% CI: 2.1-6.1) to 1.1 ± 0.2 Hz (SNX-482, n=10, 95% CI: 0.2-2.1, p=0.0036, paired Student’s t-test; Fig. 1A, B) and decreased the regularity of pacemaking (coefficient of variation of the mean interspike interval increased from 0.25 ± 0.06 (control, 95% CI: 0.15-0.39) to 0.78 ± 0.13 (SNX-482, 95% CI: 0.49-1.09, p=0.0032, paired Student’s t-test; Fig. 1B)). Slowing of firing was associated with hyperpolarization of the most negative voltage reached during the afterhyperpolarization (AHP) immediately after the spike (AHP peak), which decreased from −43.2 ± 1.3 mV (control, 95% CI: −45.8 - −40.7) to −47.0 ± 1.2 mV (SNX-482, 95% CI: −49.4 - −44.6, p=0.0005, paired Student’s t-test; Fig. 1B). Other changes in the AP waveform, which could represent indirect effects from the slowing of AP frequency or result from inhibition of Cav2.3 channels, were also noted: a reduced mean AP half-width (control: 5.1 ± 0.3 ms, 95% CI: 4.4-5.8; SNX-482: 4.2 ± 0.3 ms, 95% CI: 3.7-4.8; p=0.0050, paired Student’s t-test), and a trend towards increased maximum time-derivative of voltage (control: 45.3 ± 4.9 mV/ms, 95% CI: 34.4-56.3; SNX-482: 74.3 ± 13.5 mV/ms, 95% CI: 47.0-101.5, p=0.0625, paired Student’s t-test, estimated from the phase-plane plot of Fig. 1C). The latter is likely due to the recruitment of more voltage-gated Na^+^-channels during the AP onset from the more hyperpolarized interspike membrane potential (Guarina et al., 2018; Tomagra et al., 2019).

**Figure 1.**
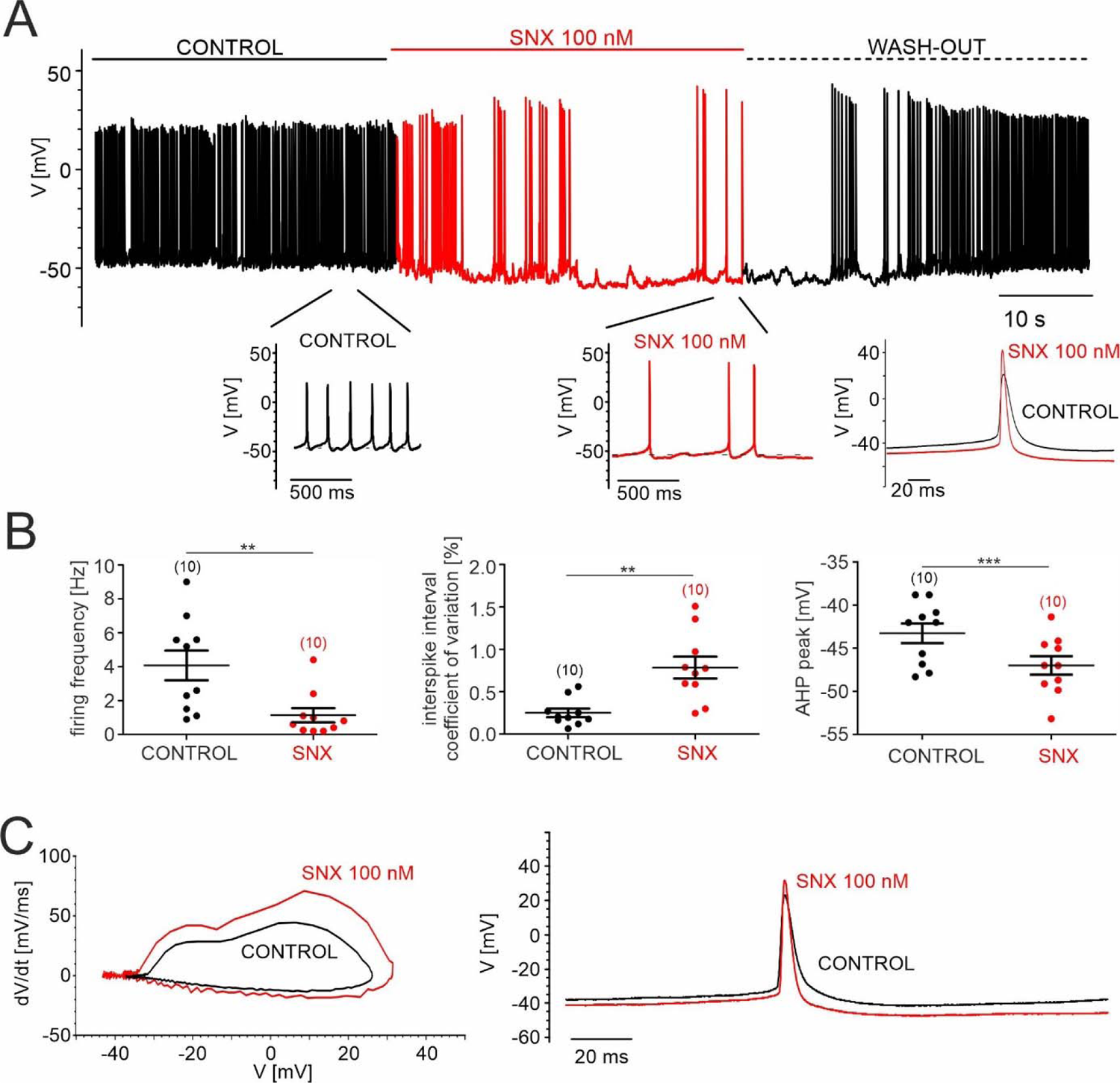
SNX-482 effects on pacemaking of cultured mouse midbrain DA neurons. **A.** Representative recording of spontaneous firing activity of cultured midbrain dopaminergic neurons before, during and after the application (wash-out) of 100 nM SNX-482. Inset (bottom right): overlay of one single AP before (control) and during the application of 100 nM SNX-482. **B.** Firing frequency [Hz], coefficient of variation of the interspike interval [%], and AHP peak [mV] before (control) and during the application of 100 nM SNX-482. Data represent the means ± SEM for the indicated number of experiments (N=3). Statistical significance was determined using paired Student’s t-test.: *** p<0.001; ** p<0.01; * p<0.05. **C.** Left panel: Phase-plane plot analysis (time derivative of voltage (dV/dt) vs. voltage (V)) before (control) and during the application of 100 nM SNX-482. Right panel: corresponding AP trace in control and in the presence of SNX-482.

We also isolated SNX-482-sensitive Cav2.3 currents in cultured DA midbrain neurons using voltage-clamp experiments. In order to exclude that part of the Ca^2+^-current inhibition by SNX-482 could be attributed to the inhibition of L-type channels, we added SNX-482 (100 nM) after the complete block of L-type currents (comprising about 25% of total Ca^2+^-current, not shown) by 3 µM isradipine (ISR). Ca^2+^-currents were elicited by consecutive depolarizing 50-ms square pulses to 0 mV from a holding potential of −70 mV every 10 s. Once stable recordings were obtained in the presence of isradipine (see representative experiments in Fig. 2A, B), 100 nM of SNX-482 were applied. Application of 100 nM SNX-482 significantly reduced non-L-type currents by 41 ± 4 % (95% CI: 33-49, n=20, N=4, paired Student’s t-test; p<0.001) (Fig. 2C) corresponding to an absolute decrease of current amplitude from 529 ± 57 pA (95% CI: 409-649) to 313 ± 33 pA (95% CI: 245-381, n=20, paired Students t-test p<0.01) (Fig. 2D). All residual I_Ca_ components were blocked by adding 2 μM Cd^2+^ to the bath solution (Fig. 2A, B).

**Figure 2.**
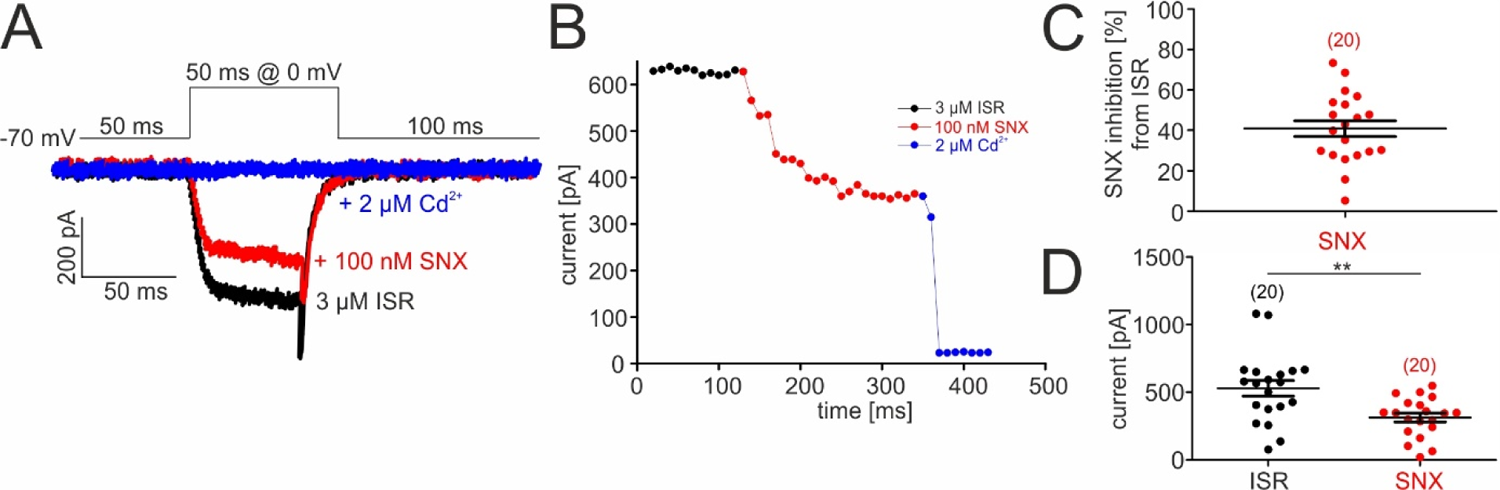
SNX-482 inhibition of non-L-type I_Ca_ in cultured midbrain DA neurons. **A.** Representative traces illustrating the inhibition of non-L-type I_Ca_ by 100 nM SNX-482 (red). Cells were initially perfused with a bath solution containing 3 μM isradipine (ISR, black). Full block was obtained using 2 μM Cd^2+^ (blue). Square pulses (50 ms) were applied to 0 mV from a holding potential of −70 mV (top) **B.** Current amplitude values plotted as a function of time. After stabilization of I_Ca_ with ISR (black circles), 100 nM SNX-482 was applied. The remaining currents was blocked by 2 μM Cd^2+^. **C.** SNX-482 inhibition expressed as % of control I_Ca_ after LTCC block using 3 μM ISR. **D.** Mean current amplitude at the end of ISR application and at the end of SNX-482 application. Data represent the means ± SEM for the indicated number of experiments (N=4). Statistical significance was determined using paired Student’s t-test: *** p<0.001; **p<0.01; *p<0.05.

Primary cultures of DA neurons are not very different from freshly dissected identified midbrain DA neurons (Puopolo et al., 2007) and thus behave differently from SN DA neurons in slices (Guzman et al., 2009). Nevertheless, our experiments clearly demonstrate that Cav2.3 channels contribute to total Ca^2+^-current and can support pacemaking in cultured mouse DA neurons. This finding requires that Cav2.3 channels must be continuously available throughout the average interspike membrane potentials of these cells (between −70 to −40 mV). However, Cav2.3 α1-subunits have originally been cloned as a low-voltage gated channel with a negative steady-state inactivation voltage range (V_0.5,inact_ −78 mV; Soong et al., 1993) with almost complete inactivation at voltages positive to −50 mV. R-type currents with such negative steady-state inactivation properties have also been described in multiple studies in both native neocortical neurons (Almog & Korngreen, 2009; Sochivko et al., 2002), neurohypophyseal terminals (Wang et al., 1999), cerebellar granule neurons (Randall & Tsien, 1997; Tottene et al., 1996) and recombinant channels co-expressed with α2δ- and various β-subunit isoforms (Jones et al., 1998; Miranda-Laferte et al., 2014; Nakashima et al., 1998; Williams et al., 1994; Yasuda et al., 2004).

To explore how Cav2.3 channels can contribute to DA neuron Ca^2+^-entry during sustained neuronal activity we expressed Cav2.3 α1-subunits together with its accessory α2δ1 and β-subunits in tsA-201 cells under near-physiological conditions. For this purpose we employed physiological extracellular Ca^2+^ (2 mM), weak intracellular Ca^2+^-buffering (0.5 mM EGTA, see methods), and used AP waveforms recorded from SN DA neurons in mouse midbrain slices (2.5 Hz) or simulated bursts as command voltages as described (Ortner et al., 2017; see methods). Moreover, we employed the Cav2.3e α1-subunit splice variant for our recordings. Among the 6 major Cav2.3 α1-subunit splice variants we only detected Cav2.3e α1 in UV laser-microdissected mouse SN DA neurons in experiments using a qualitative single-cell RT-qPCR approach (Suppl. Fig. 2A,B).

### β-subunit isoform-dependent activity of Cav2.3 channels during SN DA neuron-like regular pacemaking activity

We first employed β3 and β4 isoforms for co-expression experiments in tsA-201 cells, because Cav2 channels in the brain appear to be biochemically associated predominantly with these isoforms (Liu et al., 1996; Scott et al., 1996). When we applied the simulated SN DA neuron regular pacemaker activity (initiated from a holding potential of −89 mV), large inward currents were observed in response to single AP waveforms (Fig. 3A). Cav2.3 channels conducted I_Ca_ during the repolarization phase of the AP (I_AP_) without evidence for inward current during the interspike interval (ISI, Fig. 3A). However, I_AP_ decreased rapidly during continuous activity and almost completely disappeared after 1 (co-transfected β3) − 2 min (co-transfected β4; Fig. 3B, C). The time course of I_AP_ decrease was best fitted by a bi-exponential function (see legend to Fig. 3). Our data, therefore, suggest that Cav2.3e α1-subunits, in complex with α2δ1 and β3 or β4, cannot support substantial inward currents during continuous SN DA neuron pacemaking activity. This is in contrast to our previously published finding of stable Cav1.3 Ca^2+^-channel activity persisting under near identical experimental conditions (Cav1.3 α1/α2δ1/β3 previously published data (Ortner et al., 2017) illustrated for comparison in Fig. 3B, C). In agreement with earlier studies (Jones et al., 1998; Yasuda et al., 2004), we found that recombinant Cav2.3 channels associated with β3 and β4 inactivate at negative voltages (V_0.5,inact_ < −70 mV) with a rapid inactivation time course (≥ 50% within 50 ms, Fig. 4A-C, Table 1). These characteristics can explain the observed rapid loss of Cav2.3 activity during regular pacemaking.

**Figure 3.**
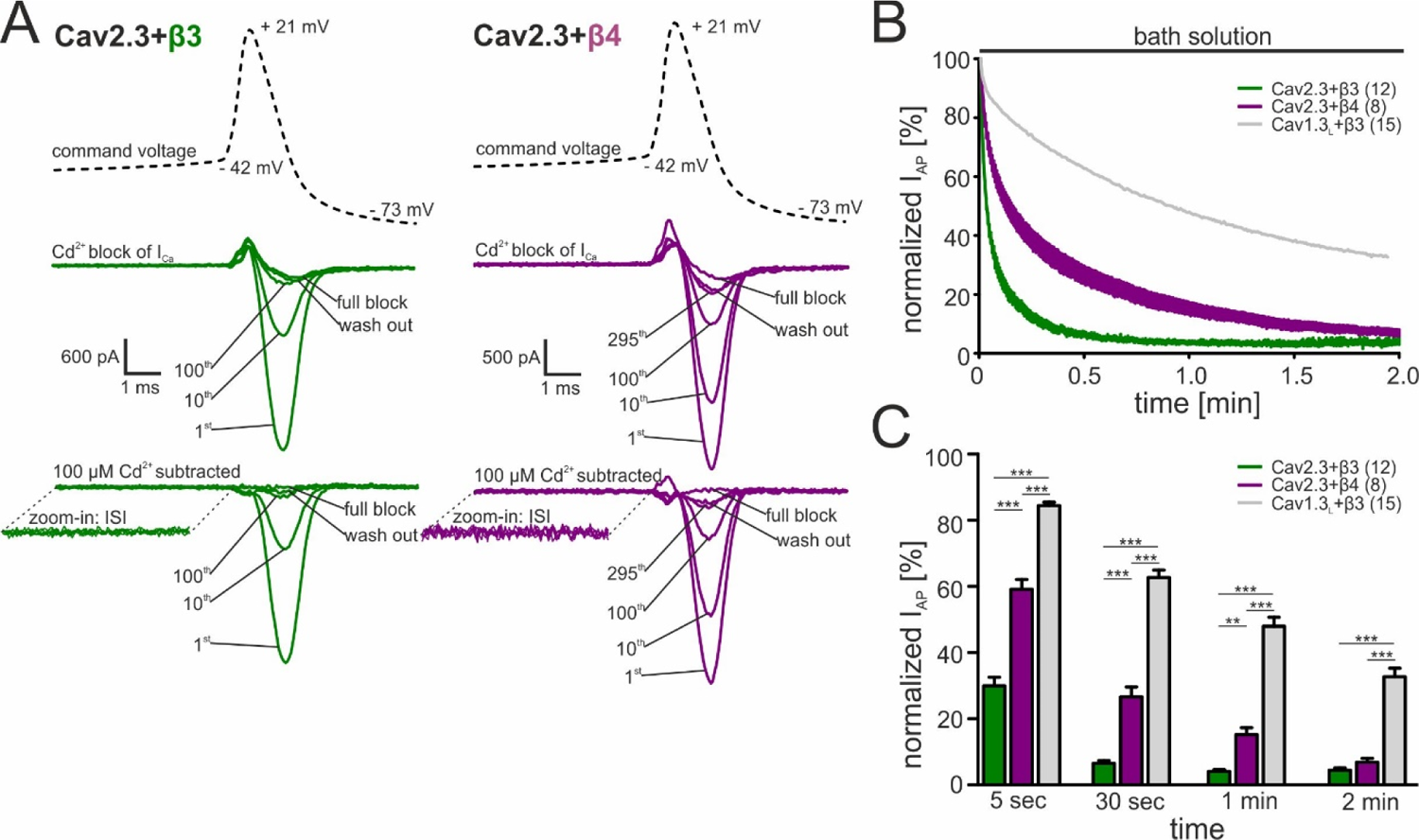
Activity-dependent inactivation of Cav2.3 channels co-transfected with β3 or β4 (and α2δ1) during simulated SN DA neuron regular pacemaking activity in tsA-201 cells. **A.** Top panel: The SN DA neuron-derived command voltage was applied with a frequency of 2.5 Hz (only a time interval around the AP-spike is shown). Middle panel: Corresponding representative Ca^2+^ current traces (2 mM charge carrier) for Cav2.3 channels co-expressed with α2δ1 and β3 (green) or β4 (purple). Cav2.3 currents were completely blocked by 100 μM Cadmium (Cd^2+^), and remaining Cd^2+^-insensitive current components were subtracted off-line (bottom panel). ISI, interspike interval. **B.** Current decay during simulated 2.5 Hz SN DA neuron pacemaking. Normalized peak inward current during APs (I_AP_) is plotted against time as mean ± SEM for the indicated number of experiments. I_AP_ amplitudes were normalized to the I_AP_ amplitude of the first AP after holding the cell at −89 mV. Cav1.3_L_ co-expressed with α2δ1 and β3 (gray, mean only) is shown for comparison (data taken from Ortner et al., 2017). The I_AP_ decay was fitted to a bi-exponential function (Cav2.3 β3: A_slow_ = 39.4 ± 0.65 %, τ_slow_ = 22.2 ± 0.15 min, A_fast_ = 54.2 ± 0.76 %, τ_fast_ = 2.86 ± 0.07 min, non-inactivating = 4.47 ± 0.13 %; β4: A_slow_ = 48.5 ± 0.26 %, τ_slow_= 90.3 ± 1.07 min, A_fast_ = 41.8 ± 0.40 %, τ_fast_ = 8.39 ± 0.16 min, non-inactivating = 5.12 ± 0.12 %). Curves represent the means ± SEM for the indicated number of experiments (N = Cav2.3/β3: 4; Cav2.3/β4: 2; Cav1.3_L_/β3: 6). **C.** Normalized I_AP_ decay after predefined time points for Cav2.3 with β3 or β4 and Cav1.3_L_ (with β3). Statistical significance was determined using one-way ANOVA followed by Bonferroni post-hoc test: *** p<0.001; ** p<0.01; * p<0.05.

**Figure 4.**
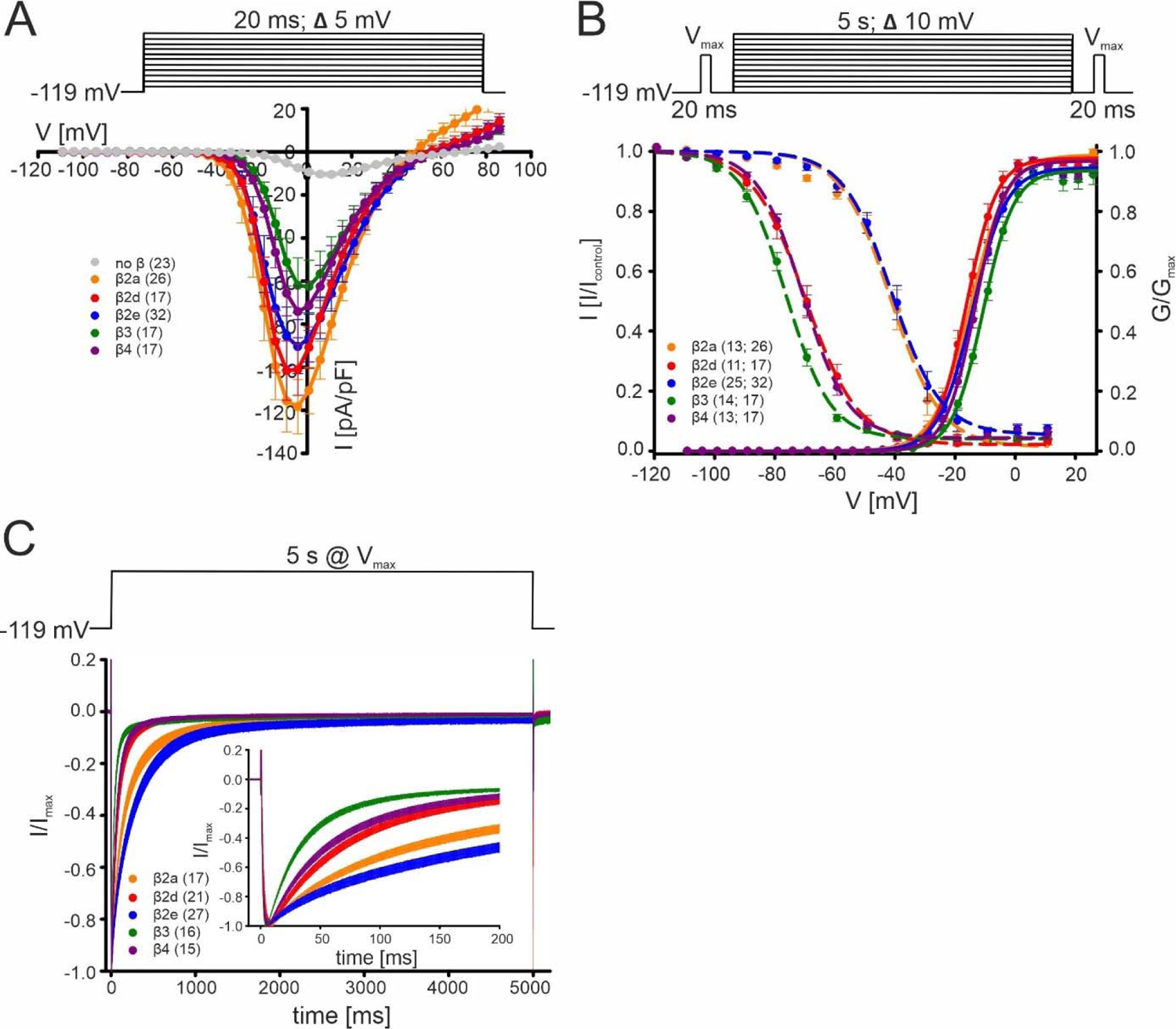
Biophysical properties of Cav2.3 channels co-transfected with different β-subunits (and α2δ1) in tsA-201 cells. **A.** Current densities (pA/pF) with or without (gray) co-transfection of indicated β-subunits. Color code and n-numbers are given in the graphs. **B.** Voltage-dependence of steady-state activation (normalized conductance G, right axis, solid lines) and inactivation (normalized I_Ca_ of test pulses, left axis, dashed lines, left n-numbers in parentheses). **C.** Inactivation time course during 5 s depolarizing pulses to V_max_ starting from a holding potential of −119 mV. Inset shows the first 200 ms of the 5 s pulse. Respective stimulation protocols are shown above each graph. The curves represent the means ± SEM for the indicated number of experiments (N = β2a: 5; β2d, β2e: 2; β3, β4, no β: 3). For statistics see Table 1. V_max_, voltage of maximal inward current.

**Table 1.**
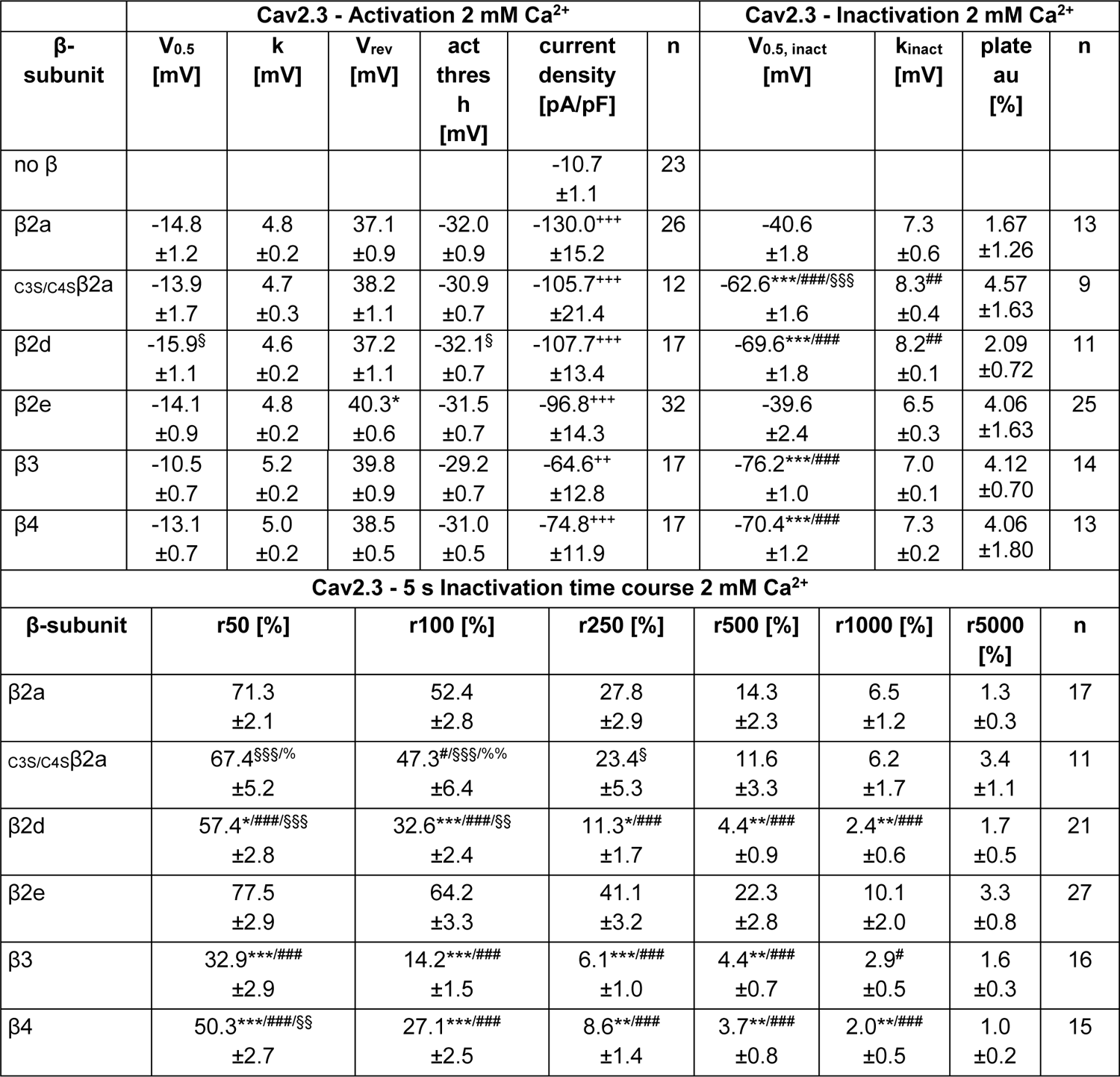
Voltage-dependence of activation and inactivation, and time course of inactivation of Cav2.3 co-transfected with α2δ1 and different β subunits in tsA-201 cells. All values are given as means ± SEM for the indicated number of experiments (N = β2a: 5; _C3S/C4S_β2a, β2d, β2e: 2; β3, β4, no β: 3). Voltage-dependence of gating: V_0.5_, Half-maximal activation voltage; k, slope factor; V_rev_, estimated reversal potential; act thresh, activation threshold; V_0.5,inact_, half-maximal inactivation voltage; k_inact_, inactivation slope factor; plateau, remaining non-inactivating current. Near physiological recording conditions (2 mM Ca^2+^, low 0.5 mM EGTA Ca^2+^ buffering) and calculation of the parameters of voltage-dependence of activation and inactivation are described in Materials and Methods. Statistical significance was determined using one-way ANOVA with Bonferroni post-hoc test (V_0.5_, V_rev_, V_0.5,inact_, k_inact_, plateau) or Kruskal-Wallis followed by Dunn’s multiple comparison test (k, act thresh, current density). Statistical significances of post-hoc tests are indicated for comparison vs. β2a (*, **, ***), vs. β2e (^#^, ^##^, ^###^), vs. β3 (^§^, ^§§^, ^§§§^) and vs. no β (^+^, ^++^, ^+++^): p<0.05, p<0.01, p<0.001. Inactivation time course: The r values represent the fraction of I_Ca_ remaining after 50, 100, 250, 500, 1000 or 5000 ms during a 5 s pulse to V_max_ (voltage of maximal inward current). Statistical significance was determined using one-way ANOVA with Bonferroni post-hoc test (r100) or Kruskal-Wallis followed by Dunn’s multiple comparison test (r50, r250, r500, r1000, r5000). Statistical significances of post hoc tests are indicated for comparison vs. β2a (*, **, ***), vs. β2e (^#^, ^##^, ^###^), vs. β3 (^§^, ^§§^, ^§§§^) or vs β4 (^%^, ^%%^, ^%%%^): p<0.05, p<0.01, p<0.001.

Since Cav2.3 channel gating properties have previously been shown to be fine-tuned by β-subunits in an isoform-dependent manner (Jones et al., 1998; Yasuda et al., 2004), we hypothesized that other β-subunits may be required to support long-lasting Cav2.3 activity during firing patterns typical for spontaneously active DA neurons. First, we tested if β-subunits are required at all for Cav2.3 activity under our experimental conditions, as Cav2.3 α1-subunits have been reported to mediate I_Ca_ even when expressed in the absence of β-subunits (Jones et al., 1998; Yasuda et al., 2004). In our experiments, all five tested β-subunits (β2a, β2d, β2e, β3, β4) caused a robust and highly significant (6-12-fold; Fig. 4A, Table 1) increase in current densities with a similar activation voltage-range. This implies that β-associated Cav2.3 channels contribute more to overall Cav2.3-mediated currents than channel complexes devoid of β-subunits and thus can be subject to differential modulation by β-subunits. Therefore, we investigated the modulation of Cav2.3 by β2-subunit isoforms, in which N-terminal alternative splicing strongly affects their modulatory effects. In particular, β2a and β2e, which contain N-termini that can anchor to the plasma membrane, can slow the inactivation time course and can affect voltage-dependence of inactivation, especially in Cav2 channels (Jones et al., 1998; Miranda-Laferte et al., 2014; Miranda-Laferte et al., 2012; Yasuda et al., 2004). In contrast, the modulatory effects of other, non-membrane anchored β2 splice variants, such as β2d, are more similar to those of β1, β3 and β4-subunits (Buraei & Yang, 2010). Indeed, also under our experimental conditions β2a and β2e subunits stabilized Cav2.3 voltage-dependent inactivation at ∼30 mV more positive potentials compared to β4 (and β3), an effect not observed for β2d subunits (Fig. 4B, Table 1). In contrast, the voltage-dependence of activation was not affected and was similar to β4 and all other tested β-subunits (Fig. 4B, Table 1). β2a and in particular β2e also significantly slowed the inactivation kinetics during prolonged depolarizations compared to all other β-subunits tested (Fig. 4C, Table 1). β3 stabilized significantly faster inactivation kinetics than β4 and β2d subunits. Due to the voltage-dependent inactivation at more depolarized potentials and the resulting overlap of the voltage-dependence of inactivation and activation (Fig. 4B), β2a and β2e subunits induce window currents at negative potentials as shown in Suppl. Fig. 3.

### β-subunit transcripts in mouse SN and VTA

The above findings prompted us to investigate if β2a and β2e are indeed expressed in SN DA neurons and could, therefore, participate in the formation of R-type currents sustained during pacemaking. We investigated β1-4 subunit expression patterns in the SN (and VTA for comparison) (Fig. 5A) dissected from brain slices of 12-14 weeks old male C57Bl/6N mice (Fig. 5C) using a standard curve-based absolute RT-qPCR assay (Schlick et al., 2010; Suppl. Table 1). In both SN and VTA tissue β4 (SN: ∼65%; VTA: ∼56%) and β2 (SN: ∼27%; VTA: ∼29%) represented the most abundant β-subunit transcripts, followed by β1 and β3 (∼5 - 7%) (Fig. 5A). Our findings are in excellent agreement (β2: 31-35%, β4: 41-45%) with cell type specific RNA sequencing data from identified midbrain DA neurons (Brichta et al., 2015; Shin, 2015).

**Figure 5.**
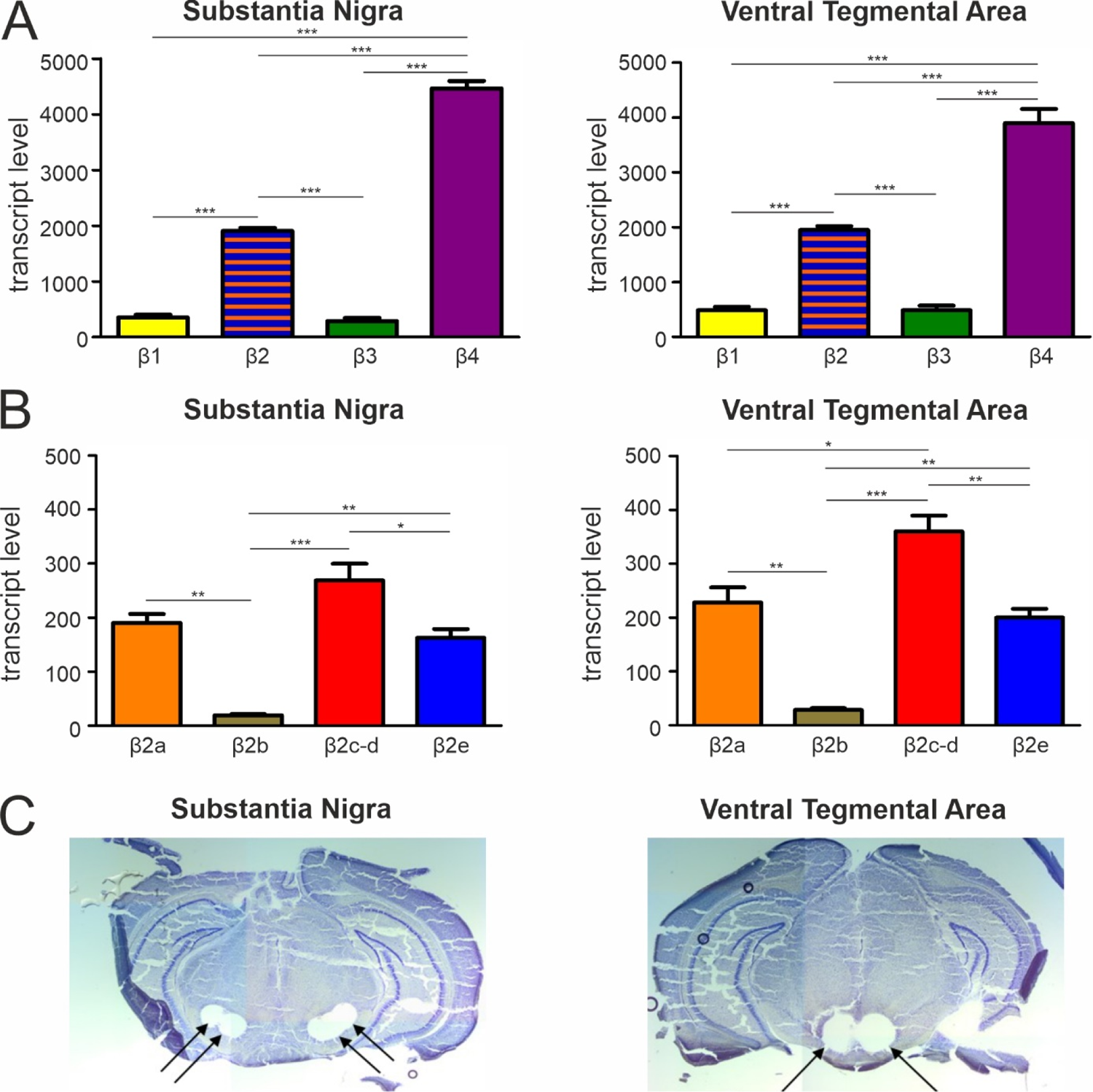
Transcript expression of various β-subunits and β2-subunit splice variants in mouse SN and VTA tissue. **A.** Expression of β1-β4 subunit transcripts in SN (n=3, N=3) (left) and VTA (n=3, N=3) (right) determined by RT-qPCR as described in Methods. **B.** Expression of β2a-β2e subunit transcripts in SN (n=3, N=3) (left) and VTA (n=3, N=3) (right). Data are shown as the mean ± SEM. Statistical significance was determined using one-way ANOVA followed by Bonferroni post-hoc test: *** p<0.001; ** p<0.01; * p<0.05. Data was normalized to *Gapdh* and *Tfrc* determined by geNorm. **C.** Example for four SN (left) and two VTA (right) tissue punches obtained for cDNA preparation with diameters of 0.5 mm each (left) or 0.8 mm each (right) from 7-8 successive 100-μm- sections between Bregma −3.00 mm and −3.80 mm, stained with Cresyl violet.

Next, we used our standard curve-based RT-qPCR assay to specifically quantify β2 N-terminal splice variant transcripts in these brain regions (Fig. 5B; for alignment of N-terminal β2 splice variants see Suppl. Fig. 1A). Assays were designed to specifically discriminate between β2a, β2b, and β2e. β2c and β2d N-termini (also present on other splice variants comprising the β2d N-terminus but with different alternative splicing in the HOOK region; Buraei & Yang, 2010) were detected by a common assay because selective primer design was difficult (see Methods). In SN and VTA, β2a (∼30%) and β2e (∼26%) transcripts together comprised about half of all tested β2-subunit splice variants, β2c and β2d-species about 42% and β2b only about 3% (Fig. 5B). Therefore, β2a and β2e together should be able to form a substantial fraction of Cav2.3 channel complexes.

We further confirmed the presence of the various N-terminal β2 splice variants in individual UV-laser microdissected mouse SN DA neurons at the mRNA level using a qualitative PCR approach (Suppl. Fig. 2C). Moreover, quantitative RNAScope analysis confirmed the expression of both, β2e and β2a variants, in identified mouse SN and VTA DA neurons with β2e more abundantly expressed compared to β2a (Suppl. Fig. 2D).

### β2 splice variant-dependent regulation of Cav2.3 activity during SN DA neuron-like regular pacemaking activity

In order to investigate if the depolarizing shifts in steady-state inactivation and slowing of inactivation kinetics by β2a and β2e subunits are sufficient to stabilize Cav2.3 currents during simulated regular pacemaking activity, we co-expressed these β-subunits with Cav2.3e channels and stimulated cells with the SN DA neuronal AP waveform as described above for β3 or β4 (Fig. 3A). In contrast to β3 or β4, I_AP_ decreased with a much slower time course in the presence of β2a or β2e with about 40% of the maximal initial I_AP_ still remaining even after 5 min of continuous activity (Fig. 6A-C). The time course of I_AP_ decrease could be fitted by a bi-exponential function predicting a steady-state reached at 32.3 ± 0.77% (n=12) of the initial I_AP_ for β2a and 25.0 ± 0.12% (n=9) for β2e (see legend to Fig. 6). Similar to co-transfected β3 or β4 subunits, β2a or β2e supported Cav2.3 I_Ca_ predominantly during the repolarization phase after the AP spike. However, in accordance with enhanced window currents at more negative voltages (Suppl. Fig. 3) these subunits also supported an inward current during the interspike interval (ISI) as evident from the first few sweeps (with the largest current amplitudes) (Fig. 6A, bottom panel, zoom-in). I_AP_ persisting after 2 min (β3, β4) or 5 min (β2a, β2e) of pacemaking was completely blocked by 100 µM Cd^2+^ and almost complete recovery from Cd^2+^-block was observed upon washout (Fig. 6B).

**Figure 6.**
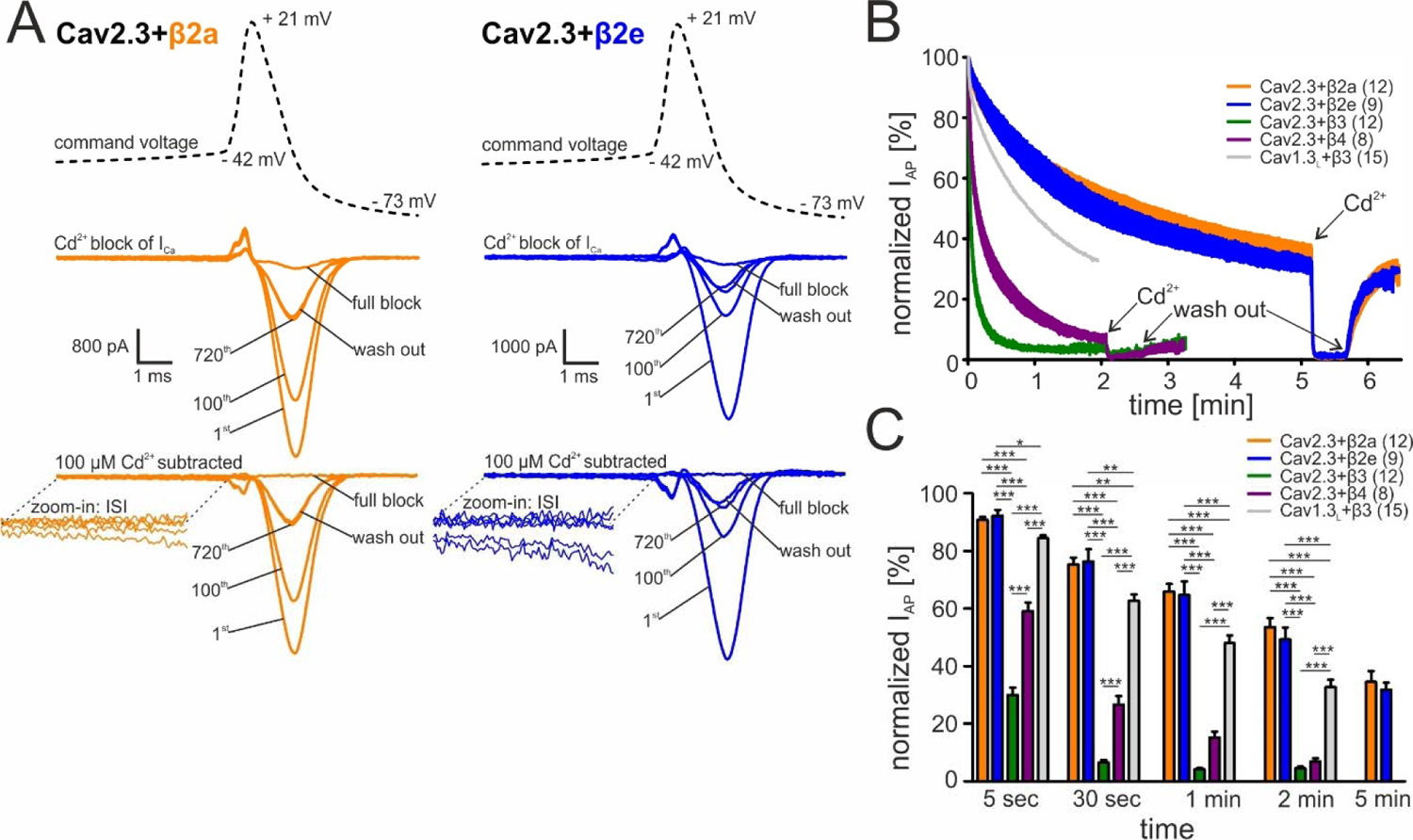
Activity-dependent inactivation of Cav2.3 channels co-transfected with β2a or β2e (and α2δ1) during simulated SN DA neuron regular pacemaking activity in tsA-201 cells. **A.** Top panel: The SN DA neuron-derived command voltage was applied with a frequency of 2.5 Hz (only a time interval around the AP-spike is shown). Middle panel: Corresponding representative Ca^2+^ current traces (2 mM charge carrier) for Cav2.3 channels co-expressed with α2δ1 and β2a (orange) or β2e (blue). Cav2.3 currents were completely blocked by 100 μM Cadmium (Cd^2+^) and remaining Cd^2+^-insensitive current components were subtracted off-line (bottom panel). **B.** Current decay (normalized I_AP_) is plotted over time as described in Fig. 3. Cav2.3 β3 (green), β4 (purple) and Cav1.3_L_/β3 data are shown for comparison (see Fig. 3). The curves represent the means ± SEM for the indicated number of experiments (N = β2a: 4; β2e: 2; β3: 4; β4: 2; Cav1.3_L_/β3: 6). The I_AP_ decay was fitted using a bi-exponential function (Cav2.3 β2a: A_slow_ = 52.6 ± 0.47 %, τ_slow_ = 299.3 ± 10.2 min, A_fast_ = 13.4 ± 0.53 %, τ_fast_ = 18.2 ± 1.47 min, non-inactivating = 32.3 ± 0.77 %; β2e: A_slow_ = 67.7 ± 0.11 %, τ_slow_ = 294.1 ± 1.77 min, A_fast_ = 7.10.0 ± 0.27 %, τ_fast_ = 16.6 ± 1.24 min, non-inactivating = 25.0 ± 0.12%). **C.** Normalized I_AP_ decay after predefined time points for Cav2.3 with co-expressed β2a, β2e, β3 or β4 and Cav1.3_L_ (with β3). Statistical significance was determined using one-way ANOVA followed by Bonferroni post-hoc test (5s, 30s, 1 min, 2 min) or unpaired Student‘s t-test (5 min): *** p<0.001; ** p<0.01; * p<0.05.

Irreversible loss of I_Ca_, a phenomenon also known as current “run-down” widely described for both native and recombinant Cavs (Kepplinger et al., 2000; Ortner et al., 2017; Schneider et al., 2018) during patch-clamp recordings, may also contribute to the current decay observed during simulated pacemaking. We, therefore, quantified the contribution of current run-down for Cav2.3 co-expressed with α2δ1 and β2a or β3 subunits first by applying short (20 ms) square pulses to V_max_ (hp −89 mV) with a frequency of 0.1 Hz (Fig. 7A). With this protocol (short pulse, long inter-sweep interval, hyperpolarized hp) activity-/voltage-dependent inactivation of Cav2.3 channels should be minimized. While β3 co-transfected Cav2.3 channels showed a time-dependent current run-down after 5 min to about 60% of the peak I_Ca_, β2a prevented the current decline during this period (Fig. 7A). To further quantify the current run-down during simulated pacemaking we interrupted the pacemaking protocol after different time periods by 20 s long pauses at −89 mV to allow channel recovery from inactivation (Fig. 7B). Thus, the percent run-down can be estimated from the non-recovering current component (Fig. 7B, horizontal dashed lines). After 30 s of pacemaking the current amplitude during the first AP after the pause was similar to I_AP_ during the initial AP with both β-subunits (β3: 96.9 ± 3.63%, 95% CI: 86.8-107.0, n=5, N=2; β2a: 107.3 ± 3.66%, 95% CI: 99.3-115.4, n=12, N=2, of the initial I_AP_). After the pause that followed another 1 min of pacemaking, 83.6 ± 7.33% (95% CI: 63.2-103.9) of the initial current was still recovered with co-transfected β3 but after 2 more minutes of pacemaking recovery decreased to 54.7 ± 6.27% (95% CI: 34.7-74.7, Fig. 7B; ∼45% run-down after 4.5 min in total, n=5). This time course is in good agreement with values obtained using the square-pulse protocol (Fig. 7A). Again, run-down was largely prevented by co-expressed β2a-subunits (Fig. 7B; ∼16% run-down after 4.5 min in total, n=12).

**Figure 7.**
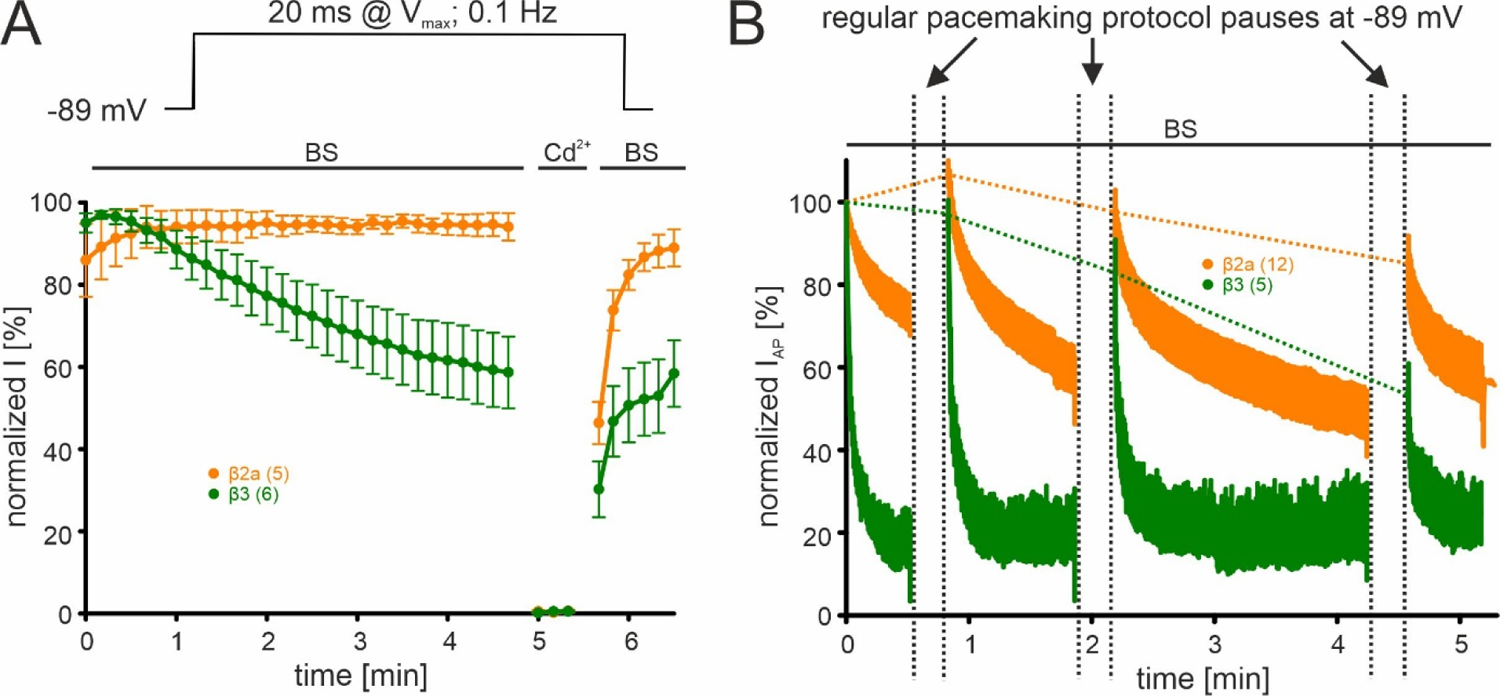
β-subunit-dependent run-down of Cav2.3 channel Ca^2+^ current in tsA-201 cells. Data for Cav2.3 co-expressed with α2δ1 and β2a (orange) or β3 (green) are shown**. A.** Run-down during a 0.1 Hz square pulse protocol (20 ms to V_max_, hp −89 mV). Currents were normalized to the I_Ca_ of the sweep with the maximal peak inward current observed during the recording. After a full block with 100 µM Cd^2+^ currents recovered to the amplitude preceding the Cd^2+^ application. **B.** Cells were held at −89 mV and then stimulated using the regular SN DA neuron pacemaking protocol for 30 s, 1 min, and 2 min each followed by 20 s long pauses (vertical dashed lines) at hyperpolarized potentials (−89 mV) to allow channel recovery from inactivation. I_AP_ of individual APs was normalized to the inward current of the first AP. The current run-down component can be estimated from the non-recovering current component (horizontal dashed lines). Traces represent means ± SEM from the indicated number of experiments (N=2). BS, bath solution

These data clearly demonstrate that the I_AP_ decrease in response to simulated SN DA neuron pacemaking (2.5 Hz) is almost completely due to the reversible accumulation of Cav2.3 channels in inactivated states, an effect partially prevented by β2a and β2e.

### Cav2.3 Ca^2+^ currents during simulated SN DA burst firing activity

In addition to regular pacemaking activity (in vitro) or irregular single spike mode (in vivo), burst firing with transient increases in intracellular Ca^2+^-load has been associated with neurodegeneration and selective neuronal vulnerability in Parkinson’s disease (Dragicevic et al., 2015; Schiemann et al., 2012). Thus, we investigated to which extent Cav2.3 Ca^2+^-channels can contribute to Ca^2+^-entry during bursts and after post-burst hyperpolarizing pauses of SN DA neurons. After reaching steady-state I_AP_ during simulated SN DA neuron pacemaking (β4: 1-2 min, β2a/β2e: 5-6 min, see also Fig. 6B) we applied a computer modeled typical three-spike burst, followed by a 1.5 s long afterhyperpolarization-induced pause at more negative potentials as a command voltage as previously described (Ortner et al., 2017) (Fig. 8A). First, we quantified to which extent total burst Ca^2+^-charge, i.e. I_Ca_ integrated over the duration of the burst, changes as compared to total Ca^2+^-charge during the same duration of a single AP in steady-state (calculated as the mean of 3 APs preceding the burst). Integrated I_Ca_ during the burst was 4-6-fold higher than the mean integrated I_Ca_ during a steady-state AP with all co-expressed β-subunits (β2a, β2e, β4) (Fig. 8A, B). However, it has to be considered that with co-expressed β4 only ∼6% of the initial I_AP_ remained in steady-state during regular pacemaking. Therefore, this relative increase will cause a much smaller rise in absolute Ca^2+^ charge compared to β2a/β2e-associated Cav2.3 where ∼40% of I_AP_ persisted in steady-state (Fig. 6B).

**Figure 8.**
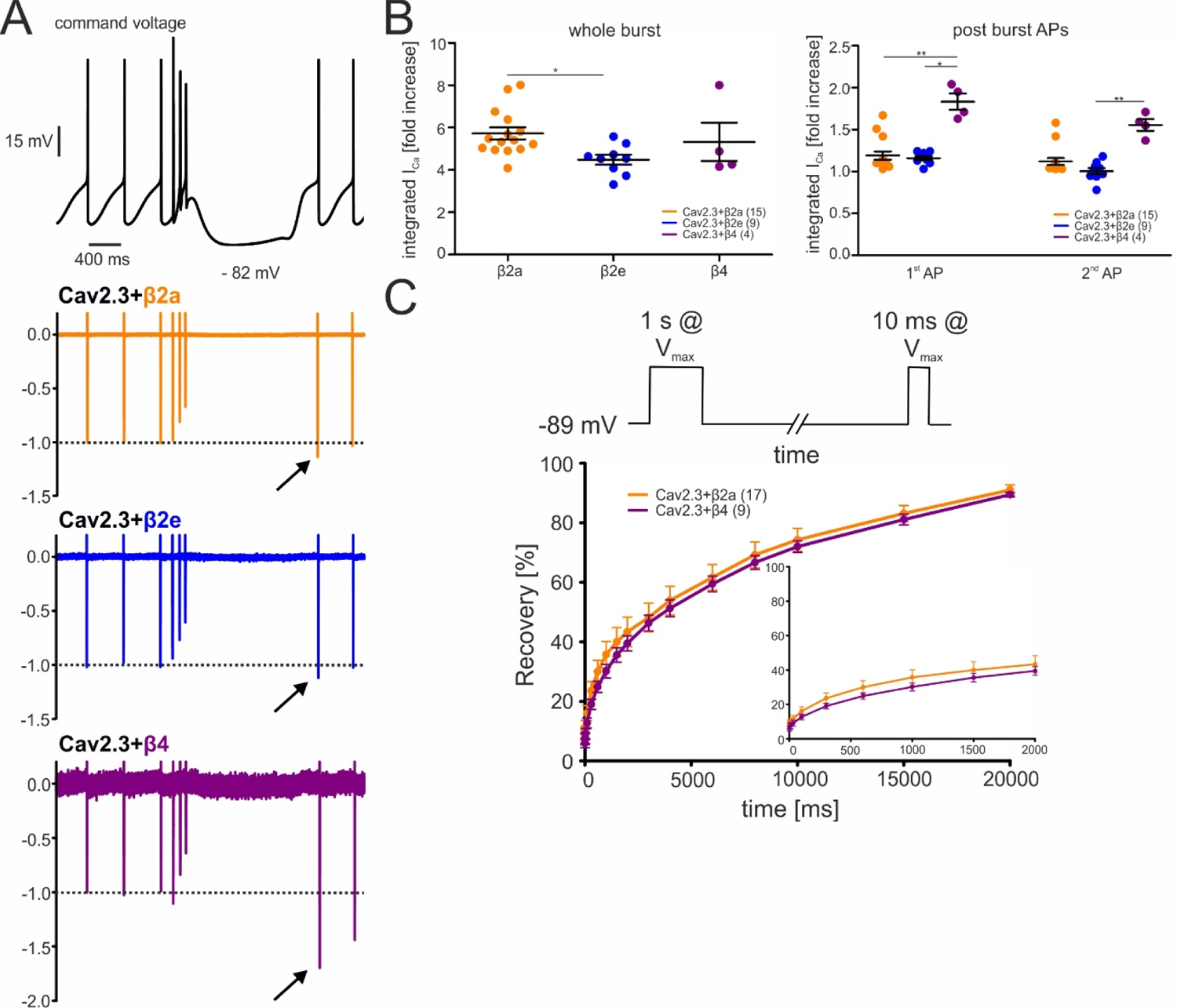
Effects of different β-subunits on Cav2.3 currents during a simulated SN DA neuron three-spike burst and post-burst APs in tsA-201 cells. The burst command voltage was elicited after ∼5-6 min (β2a, β2e) or ∼1-2 min (β4) of regular pacemaking to reach steady-state I_AP_ (β2a and β2e: ∼30% of the initial I_AP_, β4: ∼6% of the initial I_AP_, see Fig. 6). **A.** Normalized current responses of Cav2.3 channels co-expressed with β2a, β2e or β4 subunits (and α2δ1) induced by a command voltage (top panel) simulating a typical three-spike burst followed by a hyperpolarization phase at hyperpolarized potentials (lowest voltage: −82 mV) for 1.5 seconds. Remaining Cd^2+^-insensitive current components (100 µM Cd^2+^) were subtracted off-line to extract pure Cav2.3 mediated I_Ca_. One of at least four experiments with similar results is shown. **B.** The integrated I_Ca_ during a single AP before the burst (obtained as the mean of the three preceding APs) was set to 100 % and compared with I_Ca_ during the three-spike burst integrated over the time period equivalent to one AP (left) or the first APs after the pause (right). All investigated β-subunits resulted in increased integrated I_Ca_ during the burst. Data represent the means ± SEM for the indicated number of experiments (N = β2a: 4; β2e: 2; β4: 2). Statistical significance was determined using one-way ANOVA followed by Bonferroni post-test (whole burst) or Kruskal-Wallis followed by Dunn’s multiple comparison test (post-burst APs): *** p<0.001; ** p<0.01; * p<0.05. **C.** Square-pulse protocol (top) used to determine recovery from inactivation after the indicated time intervals for β2a and β4-associated Cav2.3 channels (see Methods for details). Data represent the means ± SEM for the indicated number of experiments (N=3). For statistics see Table 2.

**Table 2.** Recovery from inactivation of Cav2.3 channels co-transfected with either β2a or β4 in combination with α2δ1 in tsA-201 cells. All values are presented as the mean ± SEM for the indicated number of experiments (N=3). The r values represent the fraction of recovered I_Ca_ after 100, 1500, 4000 or 10000 ms at −89 mV between depolarizations to V_max_. No statistical significance was observed (unpaired Student’s t-test).

| Cav2.3 - recovery from inactivation 2 mM Ca<sup>2+</sup> |  |  |  |  |  |
| --- | --- | --- | --- | --- | --- |
| β-subunit | r100 [%] | r1500 [%] | r4000 [%] | r10000 [%] | n |
| β2a | 15.7<br>±2.4 | 39.3<br>±4.6 | 53.4<br>±4.5 | 74.2<br>±3.6 | 17 |
| β4 | 12.7<br>±1.7 | 35.6<br>±2.4 | 51.3<br>±2.8 | 72.0<br>±1.9 | 9 |

We also investigated if post-burst afterhyperpolarizations would allow Cav2.3 channels to recover from inactivation and thus mediate increased Ca^2+^-entry during the first APs after the burst. We first determined the recovery from inactivation of Cav2.3 channels co-expressed with α2δ1 and β4 or β2a using a square-pulse protocol (Fig. 8C; 1-s conditioning prepulse to V_max_ to inactivate Cav2.3 channels followed by a 10 ms step to V_max_ after different time periods at −89 mV). About 30% of currents recovered under these experimental conditions after 1.5 seconds at −89 mV with both co-expressed β4 and β2a (∼6-fold increase of the remaining I_Ca_ after 1s at V_max_). In contrast, recovery during the hyperpolarizing pause after the burst of the AP protocol was much less pronounced (β4) or absent (β2a, β2e) (Fig. 8B). This may be due to the different pulse protocols inducing channel inactivation over different time periods. This may stabilize inactivated states with different recovery times. Taken together, these data predict that β2a- and β2e-associated Cav2.3 channels can contribute to enhanced Ca^2+^-entry during brief burst activity but not during post-burst APs.

Since palmitoylation is reversible and regulated in an activity-dependent localized manner (Bijlmakers & Marsh, 2003; Matt et al., 2019), we also investigated the contribution of palmitoylation of β2a for Cav2.3e modulation under our experimental conditions. To mimic the de-palmitoylated form, we replaced the two N-terminal cysteines to serines (_C3S/C4S_β2a) which prevents plasma membrane anchoring of β2a (Gebhart et al., 2010; Qin et al., 1998). As shown in Suppl. Fig. 4, _C3S/C4S_β2a significantly shifted V_0.5,inact_ of Cav2.3 to more positive voltages as compared to β3 but to a much smaller extent (< 14 mV) than β2a (+35 mV) (Table 1). Due to this prominent role of palmitoylation on the V_0.5,inact_ of Cav2.3 channels, the palmitoylation state of β2a should allow further fine-tuning of non-inactivating current components of Cav2.3 channels in SN DA neurons. As illustrated in Suppl. Fig. 4, the effects of β2a palmitoylation on the inactivation kinetics and inactivation voltage of Cav1.3 L-type channels (Suppl. Fig. 4A,C) were different from Cav2.3, suggesting that palmitoylation/depalmitoylation events would regulate Ca^2+^ channel function in a subtype-selective manner.

### Steady-state activation and inactivation of R-type currents in mouse SN DA neurons

We have recently shown in whole-cell patch-clamp recordings that 100 nM SNX-482 inhibit ∼30% of total Cav currents in mouse SN DA neurons (Benkert et al., 2019) when stimulated from a holding potential of −70 mV. If membrane-anchored β2-subunits stabilize more positive V_0.5,inact_ of Cav2.3 channels in SN DA neurons then the voltage-dependent inactivation of the R-type currents should allow channels to be available even at voltages positive to −40 mV. Using whole-cell patch-clamp recordings (Fig. 9), we therefore measured the steady-state inactivation of R-type I_Ca_ in identified SN DA neurons in midbrain slices. R-type current was isolated as the current remaining after preincubation of slices with selective inhibitors of Cav1 (1 µM isradipine), Cav2 (Cav2.1, Cav2.2; 1 µM ω-Conotoxin MVIIC) and Cav3 (10 µM Z941)) (Fig. 9, blue traces/symbols). The voltage-dependence for steady-state inactivation for R-type I_Ca_ was ∼5 mV more positive than for total I_Ca_ (V_0.5,inact_ −52.4 ± 1.58, 95% CI: −55.8 - −49.0 vs −47.5 ± 1.38 mV, 95% CI: −50.5 - −44.6, p<0.05; Fig. 9A, Table 3) and very similar to recombinant Cav2.3 currents expressed with α2δ1 plus β2a or β2e subunits (∼ −40 mV, Table 1). R-type current evoked from −100 mV activated at about 8 mV more positive voltages than total I_Ca_ (Fig. 9A, Table 3 for statistics). This observation is also in accordance with the positive activation voltage range of Cav2.3 channels measured in tsA-201 cells.

**Figure 9.**
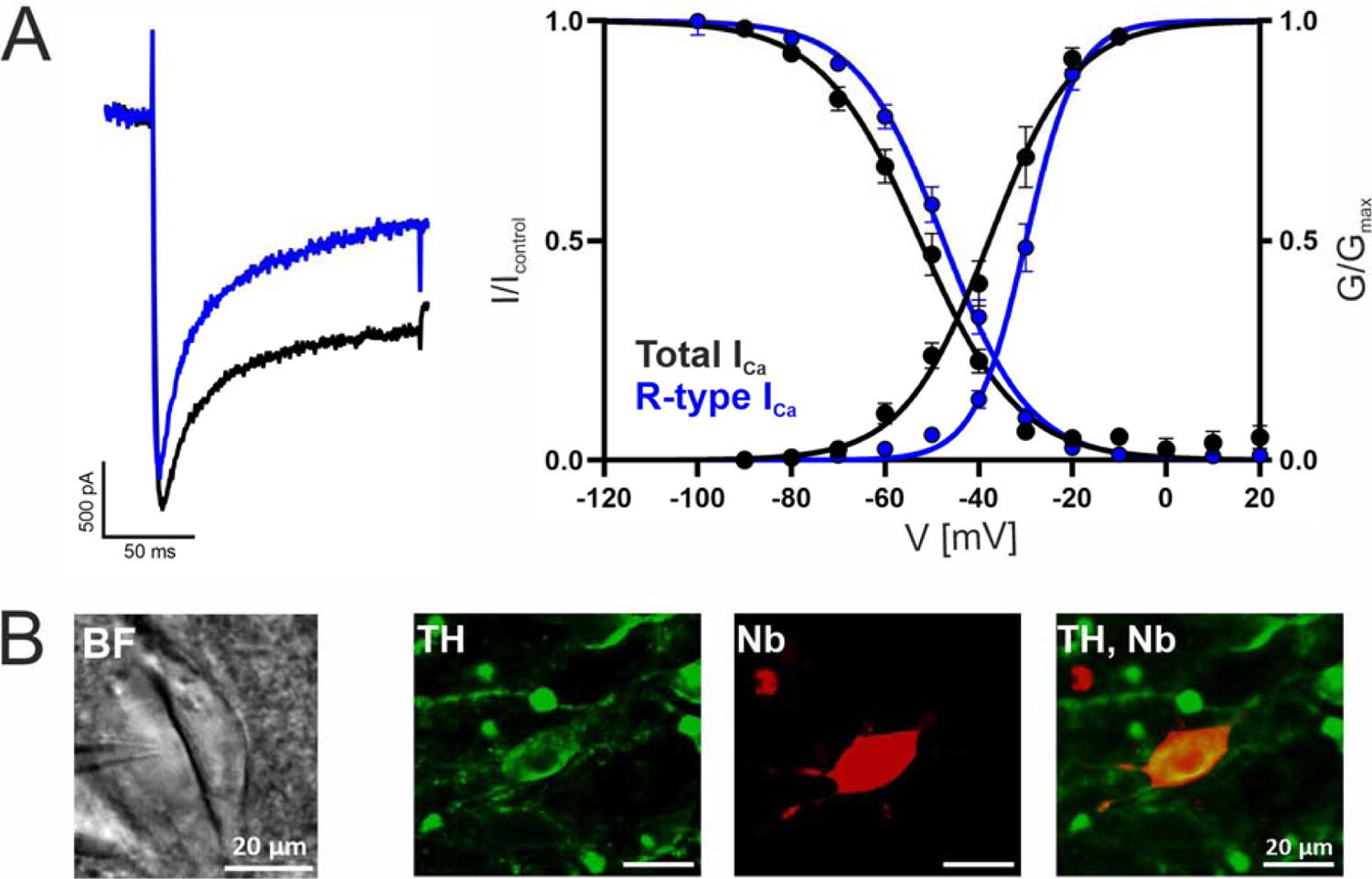
Voltage-dependence of gating of R-type currents in mouse SN DA neurons. I_Ca_ recorded in SN DA neurons without (black traces/symbols) or after preincubation (blue) of slices with a Cav channel blocker cocktail to inhibit Cav3 (10 µM Z941), Cav1 (1 µM isradipine), Cav2.1 and Cav2.2 (1 µM ω-conotoxin-MVIIC). **A.** Left panel: representative current traces of recordings of steady-state activation (at −20 mV test potential, similar amplitudes were chosen). Right panel: voltage-dependence of steady-state activation and inactivation, curve fits to the mean values (± SEM) are shown. **B.** Exemplary neuron as seen under the patch clamp microscope with patch pipette next to it (left; BF, brightfield) and a neuron after histochemical staining for tyrosine hydroxylase (TH, green) and neurobiotin (Nb, red). Detailed parameters and statistics are given in Table 3.

**Table 3.** Voltage-dependence of activation and inactivation of Ca^2+^-currents in SN DA neurons Whole cell patch-clamp experiments were performed as described in Methods in cells preincubated with a cocktail blocking T-, L-, N-, and P/Q-Ca^2+^ channels (10μM Z941, 1μM isradipine, 1μM ω-conotoxin-MVIIC) to isolate R-type currents. N, number of preparations; n, number of experiments. Voltages were not corrected for liquid junction potential (−5mV). Statistics: ****, p<0.0001; *, p<0.05 (vs. control, Mann–Whitney U test).

|  | Activation |  |  | Inactivation |  |  |
| --- | --- | --- | --- | --- | --- | --- |
| | $V_{0.5,act}$<br>[mV] | $k_{act}$<br>[mV] | n/N | $V_{0.5, inact}$<br>[mV] | $k_{inact}$<br>[mV] | n/N |
| **Control** | -37.4±1.8 | 7.2±0.58 | 16/8 | -52.4±1.58 | 9.57±0.54 | 16/8 |
| **+ T-, L-, N-, P/Q-Cav blockers** | -29.7±1.07** | 4.42±0.28*** | 19/5 | -47.5±1.38* | 8.15±0.37 | 19/5 |
| **+ 100 nM SNX-482** | -42.7±2.3 | 7.73±0.65 | 8/2 | -51.3±1.35 | 10.7±0.85 | 9/2 |

We independently confirmed the presence of SNX-482-sensitive R-type currents also using perforated patch recordings in identified SN DA neurons. From an even more positive holding potential of −60 mV, where a fraction of Cav2.3 channels must already be inactivated (Fig. 9A), still about 13% of I_Ca_ was inhibited by acute bath application of SNX-482 and inhibition was partially reversible upon washout (Suppl Fig. 5, Suppl. Table 7). These data are consistent with our finding of a contribution of β2a and/or β2e subunits to Cav2.3-mediated R-type current modulation. They also provide the first detailed analysis of Cav2.3-mediated R-type currents in identified SN DA neurons.

## Discussion

Although the modulation of Cavs by β-subunits and the characteristic gating changes induced by N-terminally membrane-anchored β2-subunit splice variants have been well described in the literature, the physiological significance of these findings remained underexplored. Here we provide strong evidence for an important role of β-subunit alternative splicing for Cav2.3 Ca^2+^-channel signaling during continuous SN DA neuron-like regular pacemaking activity. We show that only β2a and β2e stabilize Cav2.3 channel complexes with voltage-dependent inactivation properties preventing complete inactivation during the on-average depolarized membrane potentials encountered during the pacemaking cycle. This cannot only explain our finding of a substantial contribution of SNX-482-sensitive R-type currents to activity-dependent somatic Ca^2+^-oscillations in SN DA neurons in brain slices (Benkert et al., 2019) but also to pacemaking frequency in cultured DA neurons. We also provide evidence for the expression of β2a and β2e subunit splice variants in highly vulnerable SN DA and in more resistant VTA DA neurons. Together with our finding that R-type currents in SN DA neurons are available within a voltage range very similar to heterologously expressed β2a- and β2e-stabilized Cav2.3 channel complexes, our data therefore strongly suggest that these subunits are required for normal DA neuron function and may also account for the previously reported pathogenic potential of Cav2.3 Ca^2+^-channels in PD pathophysiology (Benkert et al., 2019).

We have previously shown that Cav2.3-knockout mice are protected from the selective loss of SN DA neurons in the chronic MPTP/probenecid PD model (Benkert et al., 2019). At least in this rodent model of PD, the observed protection provides strong evidence for a role of these channels in PD pathology. Pharmacological inhibition of Cav2.3 alone or together with other Cavs may, therefore, confer beneficial disease-modifying effects and a novel approach for neuroprotection in PD. The recent failure of the L-type Ca^2+^ channel blocker isradipine in the STEADY-PD trial (Parkinson Study Group, 2020) to prevent disease progression in early PD patients indicates that inhibition of L-type channels alone may not be sufficient for neuroprotection. Our previous preclinical findings identifying Cav2.3 channels as novel drug targets for neuroprotective PD-therapy are now supported by demonstrating R-type-mediated I_Ca_ in identified SN DA neurons with voltage-dependent gating properties likely stabilized by Cav2.3 association with β2a and/or β2e. Therefore, the inhibition of Cav2.3 in addition to Cav1.3 may be required for clinically meaningful neuroprotection. However, Cav2.3-mediated R-type currents are notoriously drug-resistant (Schneider et al., 2013), and so far no selective potent small-molecule Cav2.3/R-type blocker has been reported.

Our findings reported here now provide the rationale for exploring novel neuroprotective strategies based on Cav2.3 channel inhibition. These could take advantage of the strong dependence of continuous Cav2.3 channel activity on gating properties such as those stabilized by β2a and/or β2e. Rather than inhibiting Ca^2+^-entry through the pore-forming α1-subunit, such a strategy could aim at reducing its contribution to the R-type current component persisting during continuous activity in SN DA neurons by interfering with the association of β2a and β2e subunits. Even if other β-subunits would replace them in the channel complex and ensure its stable expression, our data suggest that they would not be able to shift the steady-state inactivation voltage into the operating voltage range of SN DA neurons like β2a and β2e. Such an approach appears realistic due to the availability of novel genetically encoded Ca^2+^-channel inhibitors for a cell-type-specific gene therapeutic intervention. One such approach (CaVablator) has elegantly been used to specifically target Ca^2+^-channel β-subunits for degradation by fusing β-specific nanobodies with the catalytic HECT domain of Nedd4L, an E3 ubiquitin ligase (Morgenstern et al., 2019). This strongly reduces Cav1- and Cav2-mediated Ca^2+^-currents in different cell types. At present, it is unclear to which extent other high-voltage activated Cav2 channels (Cav2.1, Cav2.2) also contribute to the high vulnerability of SN DA neurons. However, our recent quantitative RNAScope analyses in mature SN DA neurons (Benkert et al., 2019), clearly demonstrate that Cav2.3 α1-subunit (*Cacna1e*) transcripts are the most abundant α1-subunit expressed in these cells, in excellent agreement with cell type-specific RNAseq data of identified midbrain dopaminergic neurons (Brichta et al., 2015).

We also show that in cultured mouse DA neurons, Cav2.3 channels can contribute to pacemaking. SNX-482 is also a potent blocker of Kv4.3 channels (Kimm & Bean, 2014) underlying A-type K^+^-currents (I_A_). 60 nM nearly fully block reconstituted Kv4.3 currents in HEK cells (IC_50_ ∼ 3 nM). Therefore, one can argue that in current-clamp recordings, 50-300 nM SNX-482 could alter pacemaking or the AP shape by effectively blocking Kv4.3 channels. However, our data on spontaneously firing DA neurons show very clearly that 100 nM SNX-482 causes a shortening and a reduced frequency of spontaneous APs, while a block of I_A_ channels typically induces a broadening of APs in several neuronal preparations (Kim et al., 2005) and an increased frequency in DA neurons (Liss et al., 2001). Therefore, the observed changes induced by SNX-482 strongly argue for a role of non-inactivating Cav2.3-mediated R-type currents for pacemaking in DA neurons in culture.

A limitation of our work is that our experiments do not provide direct proof for a role of β2-subunit splice variants for R-type current modulation. Therefore, a similar role of other posttranslational modifications of Cav2.3 channels or protein interaction partners expressed at somatodendritic locations of SN DA neurons (such as Rab3-interacting proteins at axonal sites, Kiyonaka et al., 2007; Robinson et al., 2019) cannot be excluded. Direct proof for a role of β2-subunit splice variants would require a splice-variant-specific gene knockout or siRNA-mediated knock-down of both β2a and β2e subunits in SN DA neurons, followed by isolation of Cav2.3 current components which is methodologically challenging in these neurons. Nevertheless, we provide the first example for a physiological and perhaps even pathophysiological role of β2-subunit alternative splicing emphasizing a need for further investigation in other types of neurons.

## Materials and Methods

### Animals

For quantitative real-time PCR (RT-qPCR) experiments, male C57BL/6N mice were bred in the animal facility of the Centre for Chemistry and Biomedicine (CCB) of the University of Innsbruck (approved by the Austrian Animal Experimentation Ethics Board). For electrophysiological experiments of cultured midbrain DA neurons, C57Bl/6 TH-GFP mice (Matsushita et al., 2002; Sawamoto et al., 2001) were kept heterozygous via breeding them with C57Bl/6 mice (in accordance with the European Community’s Council Directive 2010/63/UE and approved by the Italian Ministry of Health and the Local Organism responsible for animal welfare at the University of Torino; authorization DGSAF 0011710-P-26/07/2017). All animals were housed under a 12 hours light/dark cycle with food and water ad libitum. For whole cell voltage-clamp recordings of SN DA neurons in acute brain slices, as well as single-cell RT-qPCR, juvenile male C57Bl/6 mice (PN11-13) were bred at the animal facility of Ulm University. For RNAScope analysis, adult male C57Bl/6 mice and Cav2.3 WT mice were bred at the animal facility of Ulm University. Animal procedures at the Universities of Ulm (Regierungspräsidium Tübingen, Ref: 35/9185.81-3; Reg. Nr. o.147) and Cologne (LANUV NRW, Recklinghausen, Germany (84-02.05.20.12.254) were approved by the local authorities.

### RNA isolation and cDNA synthesis for tissue RT-qPCR

Tissue was dissected after mice had been sacrificed by cervical dislocation under isoflurane (Vetflurane, Vibac UK, 1000 mg/g) anesthesia and RNA isolation and cDNA synthesis for tissue RT-qPCR was performed as described in Supplemental Methods.

### Quantitative RT-qPCR of mouse brain tissue samples

Fragments of β-subunits and β2 splice variants were amplified from mouse whole brain cDNA utilizing specific primers (Suppl. Table 1) and subcloned into the Cav1.3 8a 42 pGFP^minus^ vector after restriction enzyme digestion using SalI and HindIII. Primer sequences for β1-β4 have been described previously (Schlick et al., 2010), but additional SalI and HindIII restriction enzyme sites (underlined in Suppl. Table 1) were inserted to allow subsequent ligation of fragments into the digested vector. TaqMan gene expression assays (Thermo Fisher Scientific, Waltham, MA, USA) and custom-made TaqMan gene expression assays were designed to span exon-exon boundaries (Suppl. Table 1) as already described (Nadine J. Ortner et al., 2020).

The expression of β1, β2, β3, β4, β2a, β2b, β2c-d, and β2e was assessed using a standard curve method-based on PCR fragments of known concentration (Nadine J. Ortner et al., 2020; Schlick et al., 2010). β2c and β2d were detected by a common assay as selective primer design failed due to high sequence similarity. This assay binds at the exon-exon boundary of exons 2A and 3 of β2c and β2d and also recognizes a number of splice variants comprising the β2d N-terminus but with different alternative splicing in the HOOK region of the subunit (Buraei & Yang, 2010). Details about assay specificity are given in Suppl. Fig.1.

### cDNA constructs

For transient transfections hCav2.3e (cloned into pcDNA3, Pereverzev et al., 2002) or hCav1.3_L_ (human C-terminally long Cav1.3 splice variant; GenBank accession number EU363339) α1 subunits were transfected together with the previously described accessory subunit constructs: β3 (rat, NM_012828, Koschak et al., 2001), β2a (rat, M80545, Koschak et al., 2001), β2d (mouse, β2aN1, FM872408.1; Link et al., 2009), β2e (mouse, β2aN5, FM872407; Link et al., 2009, where β2d and β2e were kindly provided by V. Flockerzi, Saarland University, Homburg), _C3S/C4S_β2a (cysteine residues in position 3 and 4 of β2a replaced by serines, Gebhart et al., 2010) or β4 (rat, splice variant β4e, kindly provided by Dr. Bernhard Flucher, Medical University Innsbruck; Etemad et al., 2014) and α2δ1 (rabbit, NM_001082276, Koschak et al., 2001).

### Cell culture and transfection

TsA-201 cells (European Collection of Cell Culture, catalog #96121229) were cultured as described (N. J. Ortner et al., 2020) in Dulbecco’s modified Eagle’s medium (DMEM; Sigma-Aldrich, catalog #D6546) that was supplemented with 10% fetal bovine serum (FBS, Gibco, catalog #10270-106), 2 mM L-glutamine (Gibco, catalog #25030-032), penicillin (10 units/ml, Sigma, P-3032) and streptomycin (10 μg/ml, Sigma, S-6501). Cells were maintained at 37°C and 5% CO_2_ in a humidified incubator and were subjected to a splitting procedure after reaching ∼80% confluency. For splitting, cells were dissociated using 0.05% trypsin after implementing a washing step using 1 x phosphate buffered saline (PBS). TsA-201 cells were replaced and freshly thawed when they exceeded passage no. 21. For electrophysiology, cells were plated on 10-cm culture dishes and subjected to transient transfections on the following day. Cells were transiently transfected using Ca^2+^-phosphate as previously described (Ortner et al., 2014) with 3 μg of α1 subunits, 2 μg of β subunits, 2.5 μg of α2δ1 subunits and 1.5 µg of eGFP to visualize transfected cells. On the next day, cells were plated onto 35-mm culture dishes that were coated with poly-L-lysine, kept at 30°C and 5% CO_2_ and were then subjected to whole-cell patch-clamp experiments after 24-72 hours.

### Primary cell culture of midbrain DA neurons

As described in Tomagra et al., 2019, the methods for the primary culture of mesencephalic dopamine neurons from substantia nigra (SN) were adapted from Pruszak et al., 2009. Briefly, the ventral mesencephalon area was dissected from embryonic (E15) C57Bl/6 TH-GFP mice (Matsushita et al., 2002; Sawamoto et al., 2001) that were kept heterozygous via breeding them with C57Bl/6J mice. HBSS (Hank’s balanced salt solution, without CaCl_2_ and MgCl_2_), enriched with 0.18% glucose, 1% BSA, 60% papain (Worthington, Lakewood, NJ, United States), 20% DNase (Sigma-Aldrich) was stored at 4°C and used as digestion buffer. Neurons were plated at final densities of 600 cells per mm^2^ on Petri dishes. Cultured neurons were used at 8/9 days in vitro (DIV) for current-clamp and voltage-clamp experiments. Petri dishes were coated with poly-L-Lysine (0.1 mg/ml) as substrate adhesion. Cells were incubated at 37°C in a 5% CO_2_ atmosphere, with Neurobasal Medium containing 1% pen-strep, 1% ultra-glutamine, 2% B-27, and 2.5% FBS dialyzed (pH 7.4) (as previously described in Tomagra et al., 2019).

### Whole-cell patch-clamp recordings in tsA-201 cells

For whole-cell patch-clamp recordings, patch pipettes with a resistance of 1.5-3.5 MΩ were pulled from glass capillaries (Borosilicate glass, catalog #64-0792, Harvard Apparatus, USA) using a micropipette puller (Sutter Instruments) and fire-polished with a MF-830 microforge (Narishige, Japan). Recordings were obtained in the whole-cell configuration using an Axopatch 200B amplifier (Axon Instruments, Foster City, CA), digitized (Digidata 1322A digitizer, Axon Instruments) at 50 kHz, low-pass filtered at 5 kHz and subsequently analyzed using Clampfit 10.7 Software (Molecular Devices). Linear leak and capacitive currents were subtracted online using the P/4 protocol (20ms I-V protocol) or offline using a 50 ms hyperpolarizing voltage step from −89 to −99 mV or −119 to −129 mV. All voltages were corrected for a liquid junction potential (JP) of −9 mV (Lieb et al., 2014). Compensation was applied for 70-90% of the series resistance.

The pipette internal solution for recordings of Cav2.3 contained (in mM): 144.5 CsCl, 10 HEPES, 0.5 Cs-EGTA, 1 MgCl_2_, 4 Na_2_ATP adjusted to pH 7.4 with CsOH (299 mOsm/kg). The pipette internal solution for recordings of Cav1.3 contained (in mM): 135 CsCl, 10 HEPES, 10 Cs-EGTA, 1 MgCl_2_, 4 Na_2_ATP adjusted to pH 7.4 with CsOH (275 mOsm/kg). The bath solution for recordings of Cav2.3 contained (in mM): 2 CaCl_2_, 10 HEPES, 170 Choline-Cl and 1 MgCl_2_ adjusted to pH 7.4 with CsOH. The bath solution for recordings of Cav1.3 contained (in mM): 15 CaCl_2_, 10 HEPES, 150 Choline-Cl and 1 MgCl_2_ adjusted to pH 7.4 with CsOH.

Current-voltage (I-V) relationships were obtained by applying a 20 ms long square pulse protocol to various test potentials (5 mV voltage steps) starting from a holding potential (hp) of −119 mV or −89 mV (recovery of inactivation). The resulting I-V curves were fitted to the following equation:

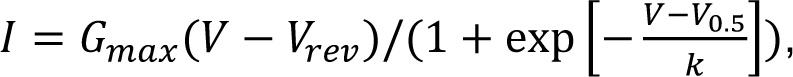

where I is the peak current amplitude, G_max_ is the maximum conductance, V is the test potential, V_rev_ is the extrapolated reversal potential, V_0.5_ is the half-maximal activation voltage, and k is the slope factor. The voltage dependence of Ca^2+^-conductance was fitted using the following Boltzmann relationship:

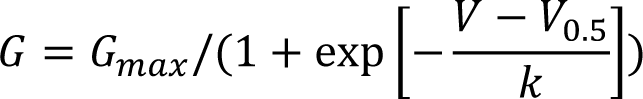

The voltage dependence of inactivation was assessed by application of 20 ms test pulses to the voltage of maximal activation (V_max_) before (I_control_) and after holding the cell at various conditioning test potentials for 5 s (30 s inter-sweep interval; 10 mV voltage steps; hp −119 mV). Inactivation was calculated as the ratio between the current amplitudes of the 20 ms test pulses. Steady-state inactivation parameters were obtained by fitting the data to a modified Boltzmann equation:

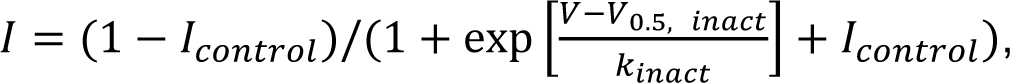

where V_0.5,inact_ is the half-maximal inactivation voltage, and k_inact_ is the inactivation slope factor.

The amount of inactivation during a 5 s depolarizing pulse from a hp of −119 mV to the V_max_ was quantified by calculating the remaining current fraction after 50, 100, 250, 500, 1000 and 5000 ms. Recovery from inactivation was determined by 10 ms test pulses to V_max_ at different time-points (in s: 0.001, 0.003, 0.01, 0.03, 0.1, 0.3, 0.6, 1, 1.5, 2, 3, 4, 6, 8, 10, 15, 20) after a 1-s conditioning pulse to V_max_ (hp −89 mV). Window current was determined by multiplying mean current densities by fractional currents form steady-state inactivation curves to obtain the fraction of available channels at a given potential as described previously (Hofer et al., 2020). The SN DA regular pacemaking command voltage protocol obtained from an identified TH^+^ SN DA neuron in a mouse brain slice (male, P12) and the SN DA burst firing protocol were generated as previously described (Ortner et al., 2017). Cells were perfused by an air pressure-driven perfusion system (BPS-8 Valve Control System, ALA Scientific Instruments) with bath solution and a flow rate of 0.6 ml/min. For Cd^2+^-block, cells were perfused with 100 µM Cd^2+^ to achieve full block, followed by wash-out with bath solution. A complete exchange of the solution around the cell was achieved within <50 ms. All experiments were performed at room temperature (∼22°C).

### Voltage- and current-clamp recordings in cultured midbrain DA neurons

Macroscopic whole-cell currents and APs were recorded using an EPC 10 USB HEKA amplifier and Patchmaster software (HEKA Elektronik GmbH) following the procedures described previously (Baldelli et al., 2005; Gavello et al., 2018). Traces were sampled at 10 kHz and filtered using a low-pass Bessel filter set at 2 kHz. Borosilicate glass pipettes (Kimble Chase life science, Vineland, NJ, USA) with a resistance of 7-8 MΩ were used. Uncompensated capacitive currents were reduced by subtracting the averaged currents in response to P/4 hyperpolarizing pulses. Off-line data analysis was performed with pClamp 10.0 software for current clamp recordings. Ca^2+^ currents were evoked by applying a single depolarization step (50 ms duration), from a holding of −70 mV to 0 mV. Fast capacitive transients due to the depolarizing pulse were minimized online by the patch-clamp analog compensation. Series resistance was compensated by 80% and monitored during the experiment.

For current-clamp experiments the pipette internal solution contained in mM: 135 gluconic acid (potassium salt: K-gluconate), 10 HEPES, 0.5 EGTA, 2 MgCl_2_, 5 NaCl, 2 ATP-Tris and 0.4 Tris-GTP (Tomagra et al., 2019). For voltage-clamp recordings the pipette internal solution contained in mM: 90 CsCl, 20 TEA-Cl, 10 EGTA, 10 glucose, 1 MgCl_2_, 4 ATP, 0.5 GTP and 15 phosphocreatine adjusted to pH 7.4. The extracellular solution for current/voltage-clamp recordings (Tyrode’s solution) contained in mM: 2 CaCl_2_, 10 HEPES, 130 NaCl, 4 KCl, 2 MgCl_2_, 10 glucose adjusted to pH 7.4. Patch-clamp experiments were performed using pClamp software (Molecular Devices, Silicon Valley, CA, United States). All experiments were performed at a temperature of 22–24°C. Data analysis was performed using Clampfit software. To study the contribution of Cav2.3 channels to the total Ca^2+^ current, cells were perfused with recording solution (containing in mM: 135 TEA, 2 CaCl_2_, 2 MgCl_2_, 10 HEPES, 10 glucose adjusted to pH 7.4) complemented with 300 nM TTX and 3 µM ISR to block voltage-dependent Na^+^ and L-type Ca^2+^ channels. SNX-482 (100 nM) was used in current- and voltage-clamp experiments. Furthermore, kynurenic acid (1 mM), 6,7-dinitroquinoxaline-2,3-dione (DNQX) (20 µM) and picrotoxin (100 µM) were present in the extracellular solution for current- and voltage-clamp experiments.

### Whole cell voltage-clamp recordings of SN DA neurons in acute brain slices

Whole-cell patch-clamp recordings were performed essentially as previously described (Benkert et al., 2019). In brief, murine (PN11-13) coronal midbrain slices were prepared in ice-cold ACSF using a VibrosliceTM (Campden Instruments). Chemicals were obtained from Sigma Aldrich unless stated otherwise. ACSF contained in mM: 125 NaCl, 25 NaHCO_3_, 2.5 KCl, 1.25 NaH_2_PO_4_, 2 CaCl_2_, 2 MgCl_2_ and 25 glucose, and was gassed with Carbogen (95% O_2_, 5% CO_2_, pH 7.4, osmolarity was 300 − 310 mOsm/kg). Slices were allowed to recover for 30 min at room temperature (22-25°C) before use for electrophysiology. Recordings were carried out in a modified ACSF solution containing in mM: 125 NaCl, 25 NaHCO_3_, 2.5 KCl, 1.25 NaH_2_PO_4_, 2.058 MgCl_2_, 1.8 CaCl_2_, 2.5 glucose, 5 CsCl, 15 tetraethylammonium, 2.5 4-aminopyridine, 600 nM TTX (Tocris), 20 µM CNQX (Tocris), 4 µM SR 95531 (Tocris) and 10 µM DL-AP5 (Tocris), pH adjusted to 7.4, osmolarity was 300 - 315 mOsm/kg. Data were digitalised with 2 kHz, and filtered with Bessel Filter 1: 10 kHz; Bessel Filter 2: 5 kHz. All recordings were performed at a bath temperature of 33°C ± 1. Patch pipettes (2.5-3.5 MΩ) were filled with internal solution containing in mM: 180 N-Methyl-D-glucamine, 40 HEPES, 0.1 EGTA, 4 MgCl_2_, 5 Na-ATP, 1 Lithium-GTP, 0.1% neurobiotin tracer (Vector Laboratories); pH was adjusted to 7.35 with H_2_SO_4_, osmolarity was 285 - 295 mOsm/kg. Neurons were filled with neurobiotin during the recording, fixed with a 4% PFA solution and stained for tyrosine hydroxylase (TH; rabbit anti-TH, 1:1000, Cat#: 657012, Merck Millipore) and neurobiotin (Streptavidin Alexa Fluor conjugate 647, 1:1000, Cat# S21374, Thermo Fisher Scientific). Only TH and neurobiotin positive cells were used for the statistical analysis.

Steady-state activation was measured by applying 150 ms depolarizing square pulses to various test potentials (10 mV increments) starting from −90 mV with a 10 sec interpulse interval. Holding potential between the pulses was −100 mV. Voltage at maximal Ca^2+^ current amplitude (V_max_) was determined during the steady-state activation recordings. Voltage-dependence of the steady-state inactivation was measured by applying a 20 ms control test pulse (from holding potential −100 mV to V_max_) followed by 5 s conditioning steps to various potentials (10 mV increments) and a subsequent 20 ms test pulse to V_max_ with a 10 sec interpulse interval. Inactivation was calculated as the ratio between the current amplitudes of the test versus control pulse. Currents were leak subtracted on-line using the P/4 subtraction. The series resistance was compensated by 60-90%. Data were not corrected for liquid junction potential (−5 mV, measured according to Neher, 1992). Midbrain slices were preincubated (bad-perfusion) at least 30 min in T-type (10 µM Z941), L-type (1 µM isradipine, ISR), N- and P/Q-type (1 µM ω-conotoxin-MVIIC) or R-type (100 nM SNX-482) Ca^2+^ channel blockers; except Z941, which was kindly obtained from T. Snutch (University of British Columbia, Canada), all Cav Blocker were from Tocris. Steady-state activation and inactivation curves were fitted as described above.

### Perforated patch recordings in SN DA neurons

Brain slice preparation was performed as described in Supplemental Methods. After preparation, brain slices were transferred to a recording chamber (∼1.5 ml volume) and initially superfused with carbogenated ACSF at a flow rate of ∼2 ml/min. During the perforation process, the electrophysiological identification of the neuron was performed in current clamp mode. Afterwards, the ACSF was exchanged for the Ca^2+^ current recording solution which contained in mM: 66.5 NaCl, 2 MgCl_2_, 3 CaCl_2_, 21 NaHCO_3_, 10 HEPES, 5 Glucose adjusted to pH 7.2 (with HCl). Sodium currents were blocked by 1 µM tetrodotoxin (TTX). Potassium currents and the hyperpolarization-activated cyclic nucleotide-gated cation current (*I*_h_) were blocked by: 40 mM TEA-Cl, 0.4 mM 4-AP, 1 µM M phrixotoxin-2 (Alomone, Cat # STP-710; Subramaniam et al., 2014) and 20 mM CsCl. Experiments were carried out at ∼28°C. Recordings were performed with an EPC10 amplifier (HEKA, Lambrecht, Germany) controlled by the software PatchMaster (version 2.32; HEKA). In parallel, data were sampled at 10 kHz with a CED 1401 using Spike2 (version 7) (both Cambridge Electronic Design, UK) and low-pass filtered at 2 kHz with a four-pole Bessel filter. The liquid junction potential between intracellular and extracellular solution was compensated (12 mV; calculated with Patcher’s Power Tools plug-in for Igor Pro 6 (Wavemetrics, Portland, OR, USA)).

Perforated patch recordings were performed using protocols modified from (Horn & Marty, 1988) and (Akaike & Harata, 1994). Electrodes with tip resistances between 2 and 4 MΩ were fashioned from borosilicate glass (0.86 mm inner diameter; 1.5 mm outer diameter; GB150-8P; Science Products) with a vertical pipette puller (PP-830; Narishige, London, UK). Patch recordings were performed with ATP and GTP free pipette solution containing (in mM): 138 Cs-methanesulfonate, 10 CsCl_2_, 2 MgCl_2_, 10 HEPES and adjusted to pH 7.2 (with CsOH). ATP and GTP were omitted from the intracellular solution to prevent uncontrolled permeabilization of the cell membrane (Lindau & Fernandez, 1986). The patch pipette was tip filled with internal solution and back filled with 0.02% tetraethylrhodamine-dextran (D3308, Invitrogen, Eugene, OR, USA) and amphotericin-containing internal solution (∼400 μg/ml; G4888; Sigma-Aldrich, Taufkirchen, Germany) to achieve perforated patch recordings. Amphotericin was dissolved in dimethyl sulfoxide (final concentration: 0.2 - 0.4%; DMSO; D8418, Sigma-Aldrich) (Rae et al., 1991), and was added to the modified pipette solution shortly before use. The used DMSO concentration had no obvious effect on the investigated neurons. During the recordings access resistance (*R*_a_) was constantly monitored and experiments were started after *R*_a_ was < 20MΩ. In the analyzed recordings *R*_a_ was comparable, did not change significantly over recording time, and was not significantly different between the distinct experimental groups. A change to the whole-cell configuration was indicated by a sudden change in *R*_a_ and diffusion of tetraethylrhodamine-dextran into the neuron. Such experiments were rejected. GABAergic and glutamatergic synaptic input was reduced by addition of 0.4 mM picrotoxin (P1675; Sigma-Aldrich), 50 µM D-AP5 (A5282; Sigma-Aldrich), and 10 µM CNQX (C127; Sigma-Aldrich) to the ACSF. For inhibition experiments, 100 nM SNX-482 (Alomone, Cat # RTS-500 dissolved in ACSF) or 10 µM nifedipine (Alomone, Cat # N-120 diluted into ACSF from a freshly prepared 10 mM stock solution in DMSO) was bath applied (in ACSF).

### Identification of β-subunit transcripts in identified SN DA and VTA DA neurons RNAScope *in situ* hybridization

*In situ* hybridization experiments were performed on fresh frozen mouse brain tissue using the RNAScope® technology (Advanced Cell Diagnostics, ACD), according to the manufacturer’s protocol under RNase-free conditions and essentially as described (Benkert et al., 2019). Briefly, 12 µm coronal cryosections were prepared (Duda et al., 2018), mounted on SuperFrost® Plus glass slides, and dried for one hour at −20°C. Directly before starting the RNAScope procedure, sections were fixed with 4% PFA for 15 min at 4°C and dehydrated using an increasing ethanol series (50%, 75%, 100%, 100%), for 5 min each. After treatment with protease IV (ACD, Cat# 322336) for 30 min at room temperature, sections were hybridized with the respective target probes for 2 h at 40°C in a HybEZ II hybridization oven (ACD). Target probe signals were amplified using the RNAScope Fluorescent Multiplex Detection Kit (ACD, Cat# 320851). All amplifier solutions were dropped on respective sections, incubated at 40°C in the HybEZ hybridization oven, and washed twice with wash buffer (ACD) between each amplification step for 2 min each. Nuclei were counterstained with DAPI ready-to-use solution (ACD, included in Kit) and slides were coverslipped with HardSet mounting medium (VectaShield, Cat# H-1400) and dried overnight. Target probes were either obtained from the library of validated probes provided by Advanced Cell Diagnostics (ACD) or self-designed in cooperation with ACD. Target probes and image acquisition are described in Supplemental Methods.

### Multiplex-nested PCR, qualitative and quantitative PCR analysis in individual laser-microdissected DA neurons

Cryosectioning, UV-laser microdissection (UV-LMD) and reverse transcription as well as multiplex-nested PCR, qualitative and quantitative PCR analysis were carried out similarly as previously described in detail (Benkert et al., 2019; Duda et al., 2018; Grundemann et al., 2011; Simons et al., 2019). All cDNA samples were precipitated prior to PCR, as described (Liss, 2002). More details are given in Supplemental Methods.

### Statistics

Data were analyzed using Clampfit 10.7 (Axon Instruments), Microsoft Excel, SigmaPlot 14.0 (Systat Software, Inc), and GraphPad Prism 5 or 7.04 software (GraphPad software, Inc). Data were analyzed by appropriate statistical testing as indicated in detail for all experiments in the text, figure and table legends. Statistical significance was set at p < 0.05. Brain slice patch-clamp data were also analyzed with FitMaster (v2×90.5, HEKA Elektronik). RNAScope and single-cell RT-qPCR data data were analyzed by Fiji (https://imagej.net/Fiji), QuantStudio^TM^ Design and Analysis Software (Applied Biosystems) and GraphPad Prism 7.04. All values are presented as mean ± SEM (95% confidence interval) for the indicated number of experiments (n) from N independent experiments (biological replications) in the text and Figures unless stated otherwise.

## Acknowledgements

This work was supported by the Austrian Science Fund (FWF, P27809, W1101, CavX-DOC 30 doc.fund) and the Tyrolean Science Fund (TWF, UNI-0404/2345), by the Italian Miur (2015FNWP34) and by the Compagnia di San Paolo (CSTO165284). We thank Dr. Veit Flockerzi for cDNA of β2 splice variants, and Jennifer Müller and Gospava Stojanovic for expert technical assistance.

## Author Contributions Statement

JS, NJO, and AS designed the study, AS performed electrophysiological recordings in tsA-201 cells, EMF dissected SN and VTA tissue, NTH prepared cDNA from mouse tissue, NTH and KV performed RT-qPCR analyses, NW performed RNAScope experiments, JB, JD performed SN DA neuron Cav2.3 splice variant analysis, JD, DS performed UV-LMD and single-cell RT-qPCR, AG, CP performed whole cell patch clamp recordings of SN DA neurons from acute mouse brain slices; GT performed electrophysiological recordings and VC and EC designed, analyzed and interpreted the experiments of AP and Cav current recordings in cultured neurons; PK and SH performed recordings in SN DA neurons in slices, JS, NJO and AS prepared the manuscript and all authors reviewed and revised the final manuscript.

## Supplement

### Supplemental Figures

**Supplemental Figure 1.**
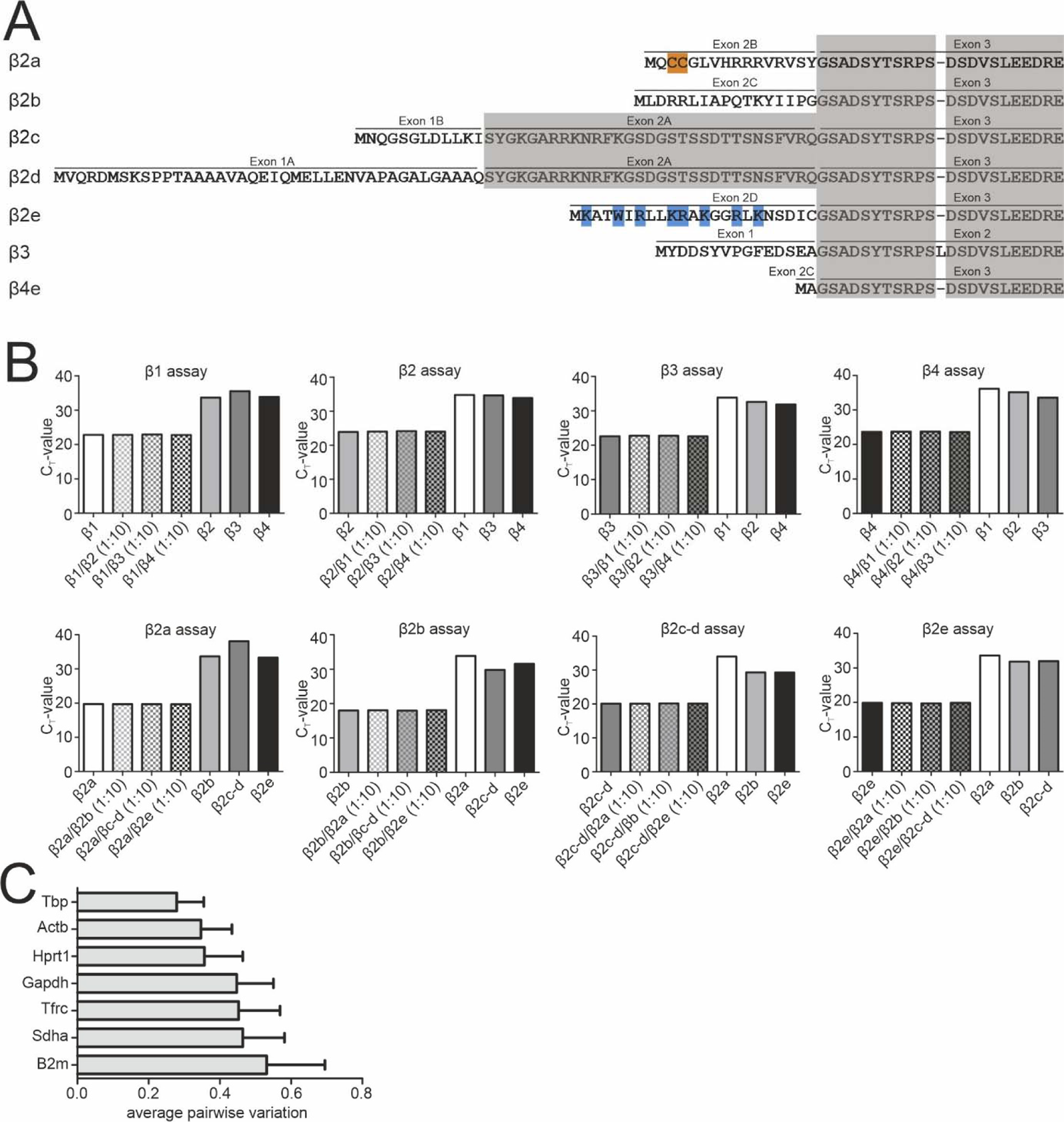
Binding specificity of β1-β4 and β2a-β2e TaqMan assays and expression stability of endogenous control genes. A. Alignment of the N-terminal amino acid sequence of the investigated β-subunits (β2a, β2d, β2e, β3 or β4). Residues responsible for membrane anchoring of β2a (palmitoylated cysteine residues are highlighted in orange) and β2e (positively charged amino acids forming a lipid binding motif are highlighted in blue) are indicated. Homologous exons are shaded in grey. Sequence accession numbers (human): β2a (NP_000715.2), β2d (NP_963887.2), β2e (NP_963864.1), β3 (NP_000716.2), β4e (NP_001307651.1). **B.** The indicated β-subunit isoforms (β1-β4; n=6) and β2 splice variants (n=3) were recognized with high specificity (low CT value) by the corresponding RT-qPCR assay even in the presence of a 10-fold higher concentration of the mismatching DNA fragments corresponding to other isoforms/splice variants (β1-4: n=2; β2 splice variants: n=1). High binding specificity was confirmed by inefficient detection (high CT value) of all non-matching DNA fragments. **C.** Average expression stability of endogenous control genes in SN and VTA tissue. All reference genes maintained comparable cDNA concentrations throughout the experiments. Data were normalized to the expression of *Gapdh* and *Tfrc* determined by geNorm. Data are given as means ± SEM (n=6, N=6).

**Supplemental Figure 2.**
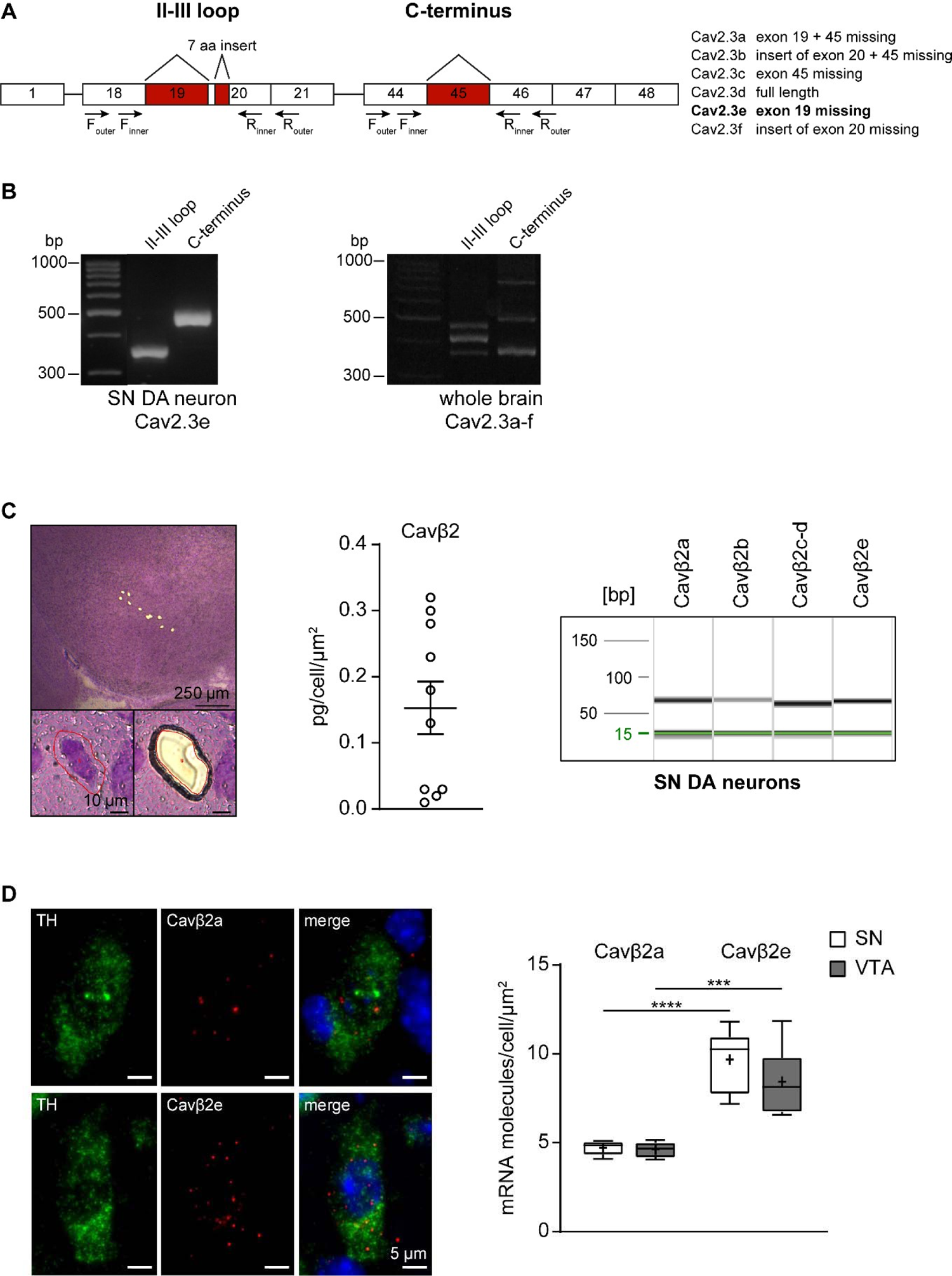
Determination of the expressed Cav2.3 splice variants in SN DA neurons. **A.** Cartoon of the exon structure of all Cav2.3 splice variants. The murine Cacna1e gene, coding for the Cav2.3 α1-subunits contains 48 exons with six major alternative splice variants (right panel). The splicing events include the variable use of exon 19, 21 nucleotides of exon 20 and exon 45. The outer and inner primer pairs chosen for identification of the different splice variants are located in the II-III loop and the C-terminus, covering the three described splicing sites. Expressed exons are shown as white boxes and splicing sites are indicated in red. **B**. Agarose gel electrophoresis image showing two PCR products (363 bp II-III loop nested PCR fragment and 498 bp C-terminus nested PCR fragment) coding for the Cav2.3e splice variant found in mouse laser-dissected SN DA neuron derived cDNA (n=40, left). In contrast, all five PCR products (363 bp, 399 bp and 420 bp in the II-III loop and 369 bp and 498 bp in the C-terminus nested PCR fragments) coding for all six major splice variants were found in whole brain tissue-derived cDNA (as positive control, right panel). **C.** Left: Overview of a cresyl violet (CV) stained coronal midbrain section of a juvenile wildtype mouse after UV-LMD of 10 SN DA neurons (scale bar: 250 μm, upper). Image of one CV stained SN DA neuron of a juvenile wildtype mouse before and after UV-LMD (scale bar: 10 μm, lower). Middle: β2-subunit transcript relative mRNA quantification by reverse transcription quantitative PCR-based in UV-LMD mouse SN DA neurons (n = 10). Right: Capillary gel electrophoresis image illustrating PCR products of β2-subunit splice variants amplified from a UV-LMD cDNA template corresponding to 2.9 cells. **D.** Left: Representative images showing β2a and β2e (red) or tyrosine hydroxylase (TH, green) RNAScope fluorescence signals (combined with nuclear DAPI staining, blue) of individual SN DA neurons from an adult wildtype mouse. Scale bar: 5 μm. Right: Absolute mRNA transcript numbers per cell for β2a or β2e in adult SN DA and VTA DA neurons of adult wildtype mice (N = 6, each), as indicated. Tukey’s boxplots are shown. Significances are indicated by asterisks: *** p < 0.001, **** p < 0.0001. For statistics see details in Suppl. Table 4.

**Supplemental Figure 3.**
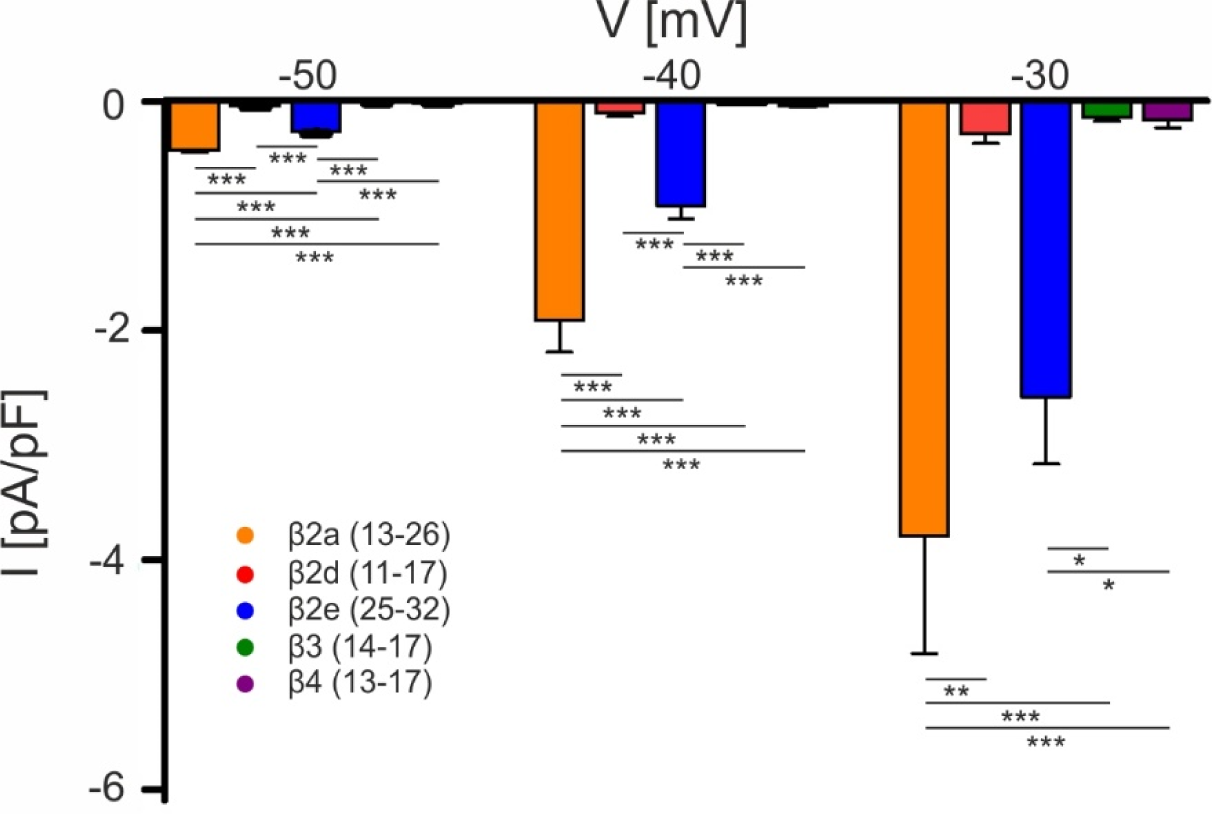
Window currents of Cav2.3 co-expressed with α2δ1 and different β-subunits. Window currents measured in the presence of the indicated β-subunits were calculated by multiplying mean current densities (pA/pF) of I-V-relationships by the fractional current inactivation from steady-state inactivation curves at the indicated voltages. Color code and n-numbers are shown in the graph. Data represent the means ± SEM for the indicated number of experiments (N = β2a: 5; β2d, β2e: 2; β3, β4: 3). Statistical significance was determined using one-way ANOVA with Bonferroni post-hoc test and is indicated: *** p<0.001; ** p<0.01; * p<0.05.

**Suppl. Figure 4.**
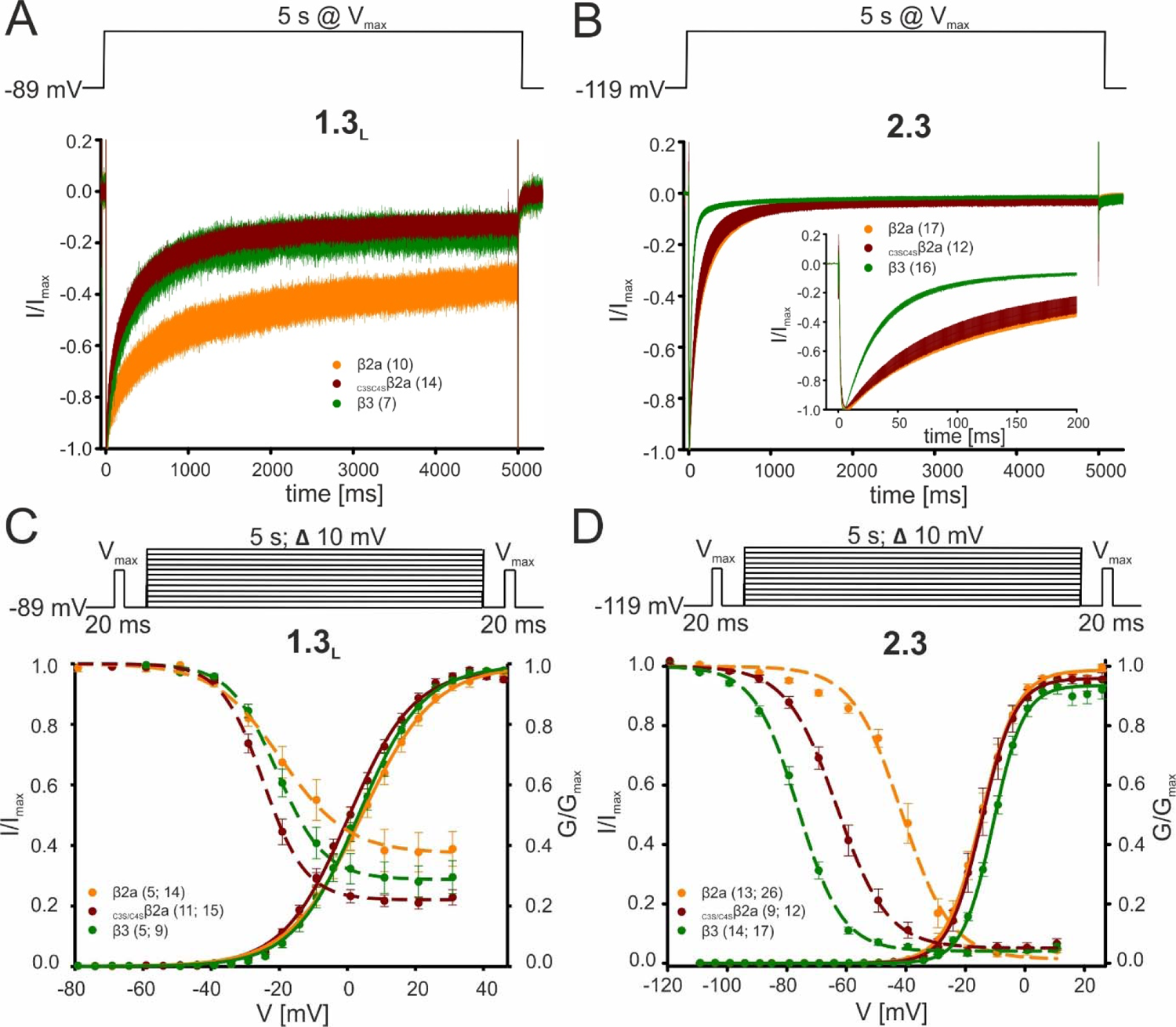
Effect of β2a palmitoylation on the biophysical properties of Cav2.3 and Cav1.3 Ca^2+^ channels in tsA-201 cells. Data are shown for Cav1.3 (C-terminally long splice variant, Bock et al., 2011) or Cav2.3 α1-subunits co-expressed with α2δ1 and β2a (orange), _C3S/C4S_β2a (red) or β3 (green). Respective command voltages are given in each panel. To disrupt palmitoylation-mediated membrane anchoring, the two N-terminal cysteines of β2a (see Suppl Fig. 1A) were replaced by serines in _C3S/C4S_β2a. **A, B.** Inactivation kinetics during a 5 s long depolarizing step to V_max_ for Cav1.3_L_ (**A**, 15 mM Ca^2+^, hp −89 mV) or Cav2.3 (**B**, 2 mM Ca^2+^, hp −119 mV). Curves represent means ± SEM for the indicated number of experiments. For statistics see Tables 1 and Suppl. Table 6. **C, D.** Voltage-dependence of activation (solid lines, normalized conductance G) and inactivation (dashed lines, normalized I_Ca_ of 20 ms test pulses) for Cav1.3_L_ (**C,** 15 mM Ca^2+^, hp-89 mV) or Cav2.3 (**D**, 2 mM Ca^2+^, hp −119 mV). Means ± SEM. For statistics see Tables 1 and Suppl. Table 6. Unlike β2a, _C3S/C4S_β2a was unable to slow the inactivation time course of L-type channels, such as Cav1.3, thus stabilizing faster inactivation similar to β3 (Suppl. Table 6, Gebhart et al., 2010). In contrast, preventing palmitoylation of β2a did not affect the inactivation time course of Cav2.3e (Table 1). Moreover, unlike observed for Cav2.3, steady-state inactivation was not significantly different for Cav1.3 co-transfected with β2a, β3, or _C3S/C4S_β2a (Suppl. Table 6). In contrast, compared to β4 and β3, _C3S/C4S_β2a significantly shifted V_0.5,inact_ of Cav2.3 to more positive voltages but to a much smaller extent than β2a (+35 mV) (Table 1).

**Suppl. Figure 5:**
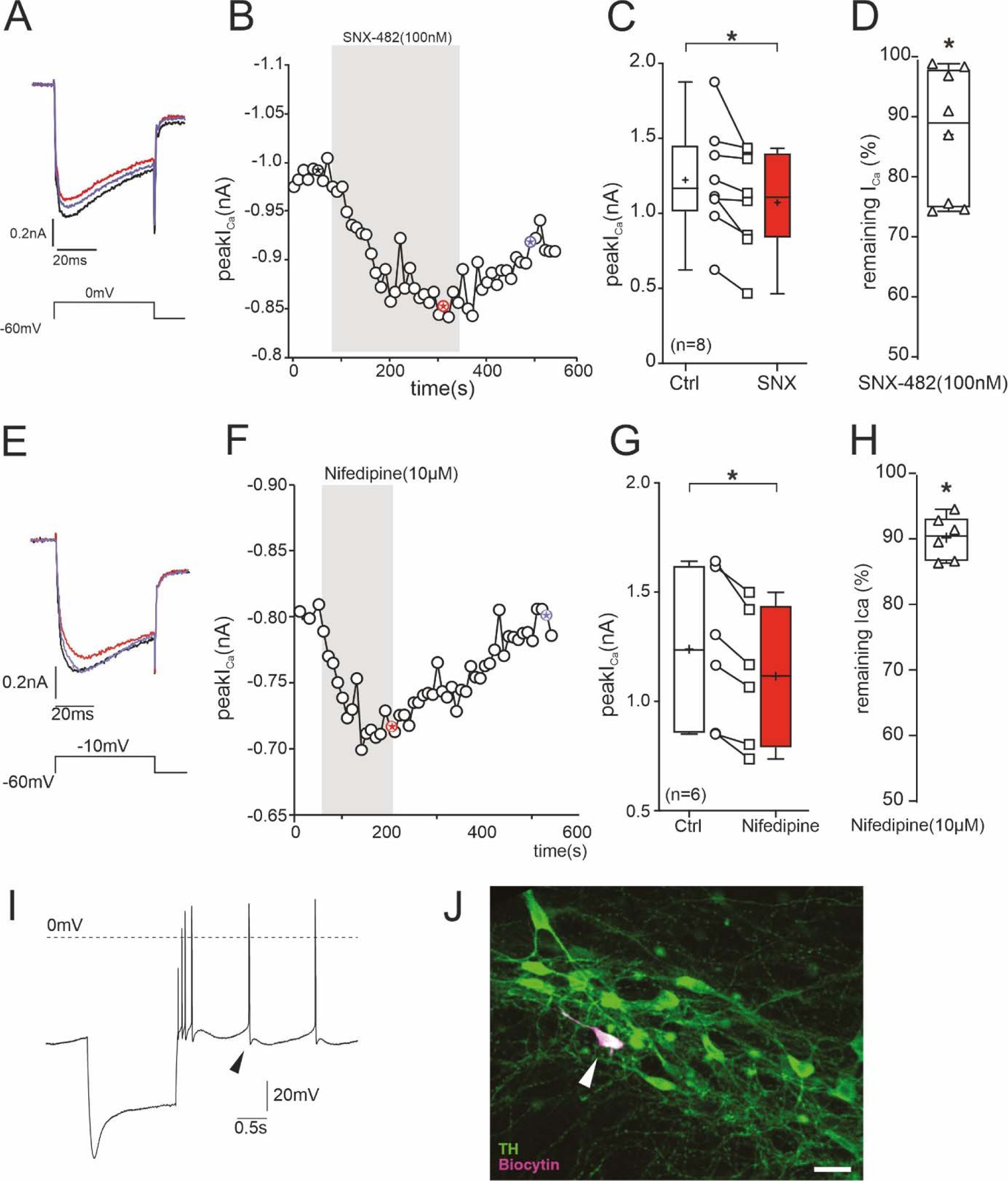
Inhibition of I_Ca_ in SN DA neurons of adult (12 weeks) mice by 100 nM SNX-482. To further prove the presence of SNX-482-sensitive R-type currents in SN DA neurons we performed perforated-patch recordings. These allow stable recordings over a long period of time and to test for reversibility of channel block by 100 nM SNX-482. Recordings were performed with the pulse protocol shown in A (holding potential −60 mV) after blocking I_A_ K^+^-currents known to be present in these cells to exclude block of I_A_ by SNX-482 (Kimm & Bean, 2014) as described previously (Benkert et al., 2019). 100 nM SNX-482 significantly reduced current to 87 ± 4% of control I_Ca_ (n=8, p=0.012, one-sample t-test, for statistics see Suppl. Table 6) and this inhibition was slowly reversible. **A.** I_Ca_ before (black), during (red) and after (blue) SNX-482 application. **B.** Example of peak I_Ca_ plotted over time during SNX-482 application. The asterisks mark the time points of the recordings shown on the left. Single depolarizing voltage steps to 0 mV were applied from a holding potential of −60 mV every 10s. **C.** Mean effect of SNX-482 on peak I_Ca_. *, p<0.05 (n=8; two-tailed paired t-test). **D.** Remaining peak I_Ca_ after SNX-482 inhibition. *, p<0.05 (two-tailed one sample t-test). Box plots: Whiskers indicate minimal and maximal values, ‘+’-sign: mean, horizontal line: median. No complete concentration-response curves were generated since higher concentrations of SNX-482 are known to also inhibit other Cav channels (Bourinet et al., 2001; Newcomb et al., 1998). Therefore, our experiments may underestimate the overall contribution of Cav2.3 channel to total I_Ca_, also because a fraction of channels must already be inactivated at the selected holding potential (−60 mV) of these experiments. **E. − H**. Under identical experimental conditions, nifedipine reduced I_Ca_ to 90 ± 1.4% of control (p=0.031, Wilcoxon-test, for statistics see Suppl. Table 6), and this inhibition was fully reversible. **E.** Perforated patch-clamp recording of I_Ca_ before (black), during (red) and after (blue) nifedipine application. **F.** Example of peak I_Ca_ plotted over time during nifedipine application. The asterisks mark the time points of the recordings shown on the left. Single depolarizing voltage steps to −10 mV were applied from a holding potential of −60 mV every 10 s. **G.** Mean effect of nifedipine on peak I_Ca_. *, p<0.05 (n=6; two-tailed paired t-test). **H.** Remaining peak I_Ca_ under nifedipine. p<0.05 (two-tailed Wilcoxon signed-rank test). Box plots: Whiskers indicate minimal and maximal values, ‘+’-sign: mean, horizontal line: median. For statistics see Suppl. Table 6. **I**. Current clamp recording of a DA neuron in the SN pars compacta to demonstrate the pre-identifying electrophysiological properties including sag potential during hyperpolarization, broad action potentials, and regular pacemaking. Arrowhead: largely reduced slow afterhyperpolarization due to partial block of K^+^ channels by Cs^+^ diffusion into the neuron during the perforation process. **J.** Post recording analysis shows co-localization of biocytin (recorded neuron; magenta) and tyrosine hydroxylase (TH, green).

## Supplemental Tables

**Supplemental Table 1.**
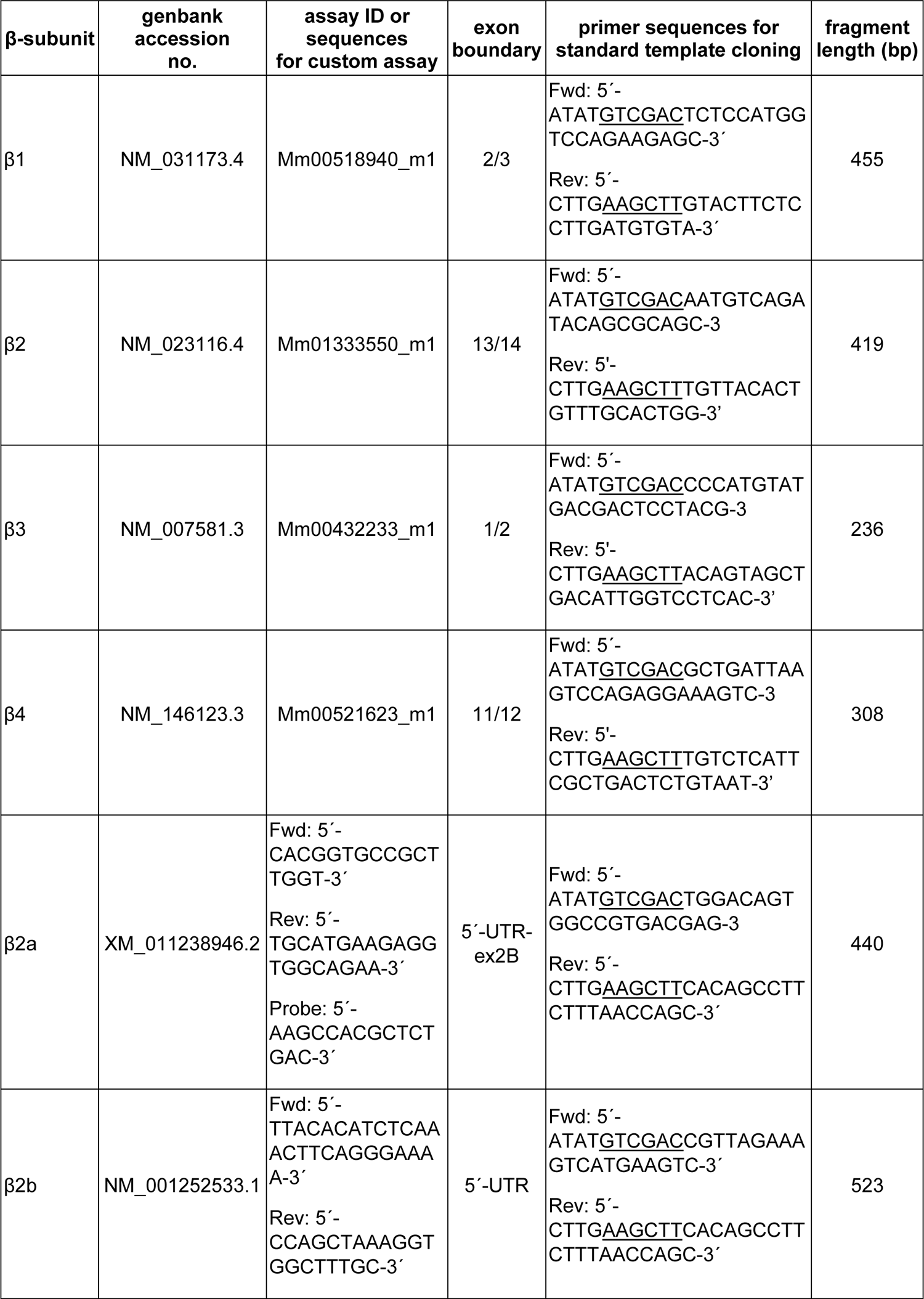
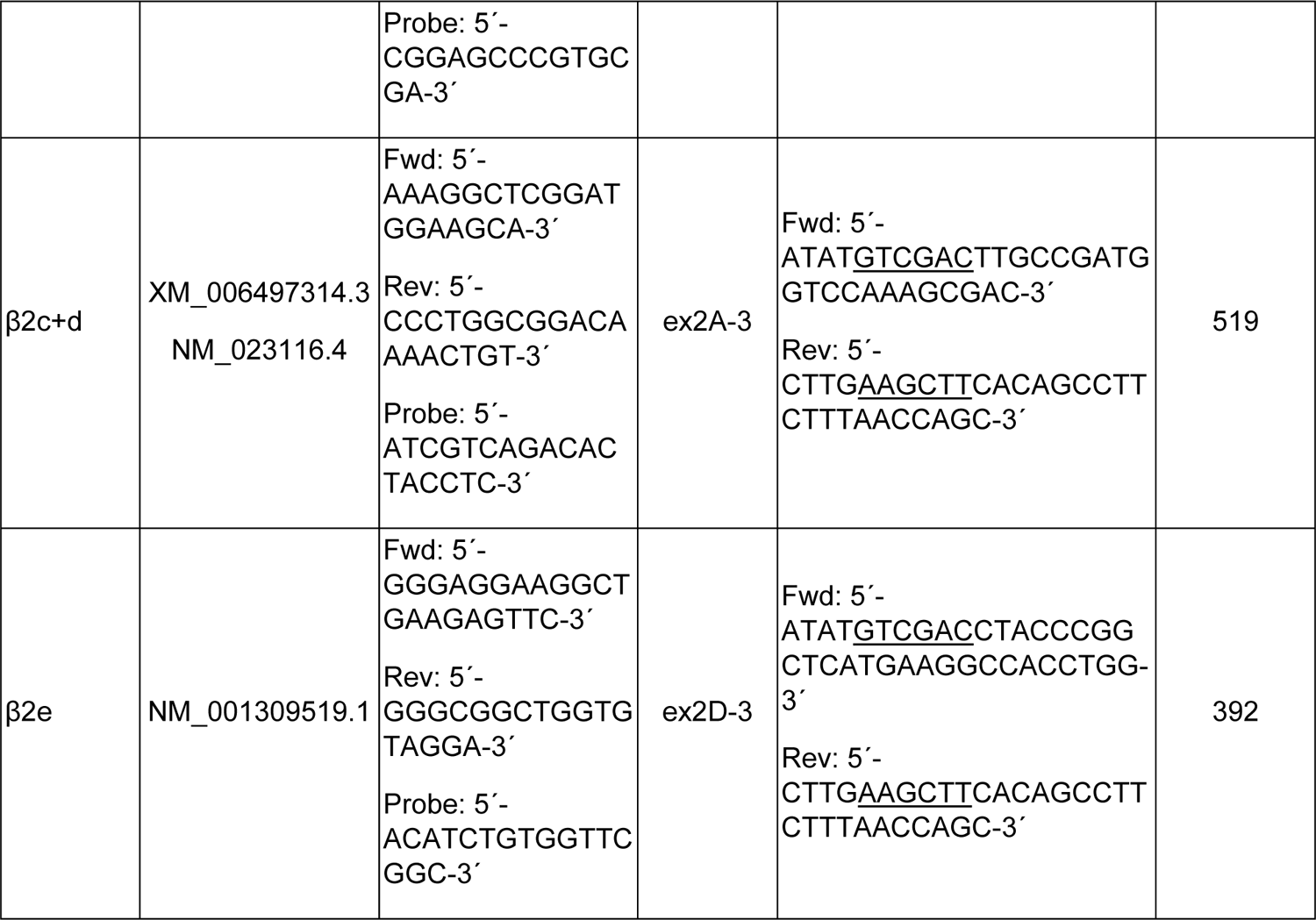
TaqMan assays for β1-4 isoforms and N-terminal β2 splice variants including cDNA specific primer sequences for standard template cloning. Additional SalI and HindIII restriction enzyme sites (see Methods) are underlined; the reverse (rev) primer used for standard template cloning including the HindIII restriction site was the same for β2a-e subunit variants; rev, reverse; fwd, forward.

**Supplemental Table 2.** Names, cellular functions and assay ID’s of endogenous control genes. For details see methods.

| gene symbol | name | cellular function | assay ID |
| --- | --- | --- | --- |
| *Actb* | beta-actin | cytoskeletal protein | Mm00607939_s1 |
| *B2m* | beta-2-microglobulin | immunity protein | Mm00437762_m1 |
| *Gapdh* | glyceraldehyde-3-phosphate dehydrogenase | oxidoreductase | Mm99999915_g1 |
| *Hprt1* | hypoxanthine phosphoribosyl-transferase 1 | transferase | Mm00446968_m1 |
| *Tbp* | tata box binding protein | transcription factor | Mm00446973_m1 |
| *Tfrc* | transferrin receptor | receptor | Mm00441941_m1 |
| *Sdha* | succinate dehydrogenase complex, subunit A | oxidoreductase | Mm01352363_m1 |

**Supplemental Table 3.**
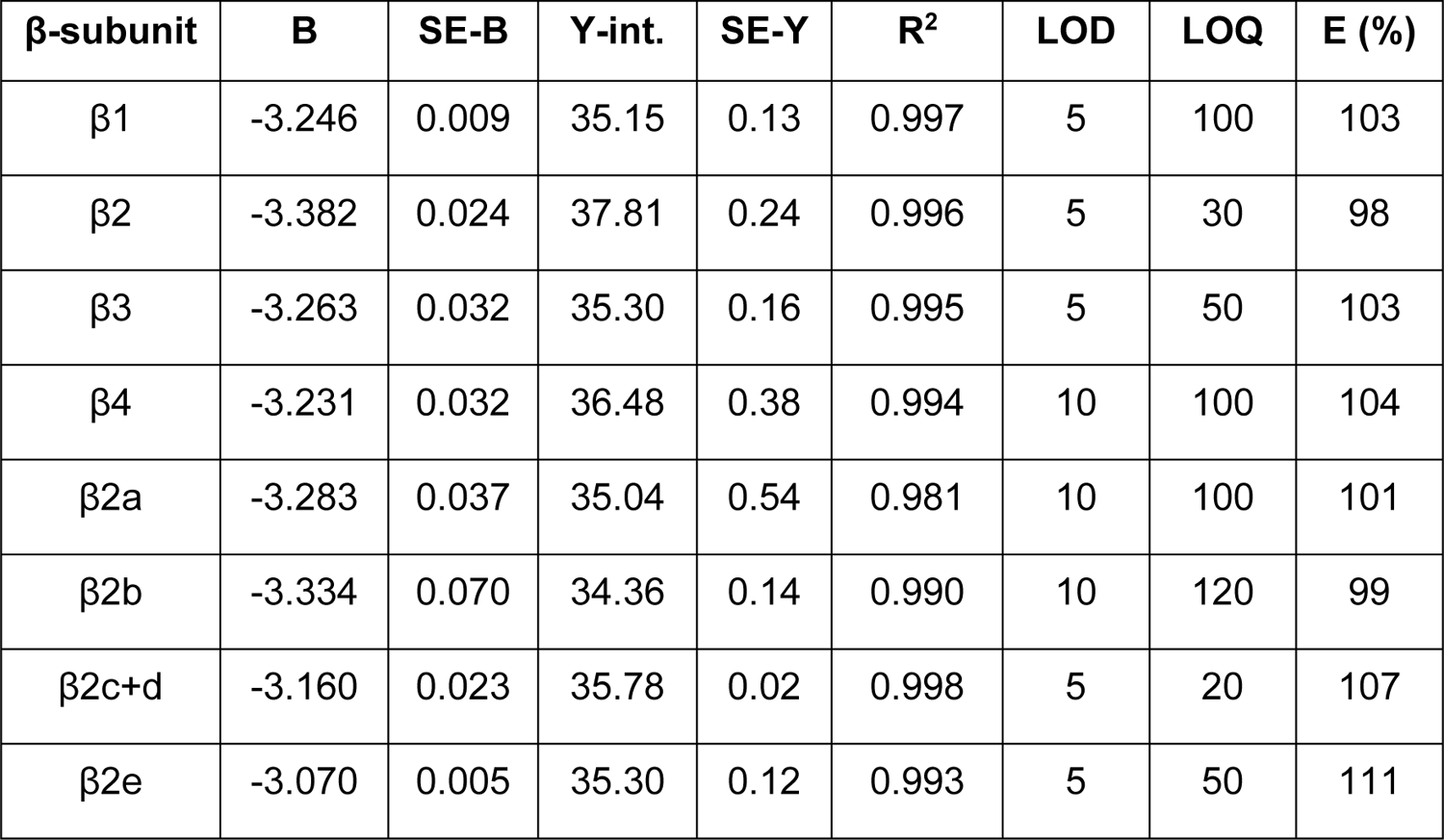
Quantitative RT-PCR standard curve parameters. SE-B,-Y, standard error of B and Y-intercept; Y-int., Y-intercept (CT value); R^2^, squared correlation coefficient; LOD, limit of detection (number of transcripts); LOQ, limit of quantification (number of transcripts); E (%), Efficiency in % (E=10-1/slope-1), 100% efficiency corresponds to a slope of −3.32.

**Supplemental Table 4:** Single cell gene expression data. Data and statistics of genes as indicated for graphs shown in Suppl Fig. 2C (middle) and D (right). n represents number of analyzed dopaminergic neurons derived from N individual mice. Significances according to two-way ANOVA followed by Tukey’s multiple comparisons test: ***, p<0.001.

|  |  |  |  |  |  |  |  |  |  |  |
| --- | --- | --- | --- | --- | --- | --- | --- | --- | --- | --- |
|  | A. mRNA expression in SN DA neurons of juvenile wildtype mice |  |  |  |  |  |  |  |  |  |
| analyzed mRNA | mean | ±SD | ±SEM | median | n/N |  |  |  |  |  |
| β2 | 0.15 | 0.13 | 0.04 | 0.16 | 10/1 |  |  |  |  |  |
|  | B. number of mRNA molecules in DA neurons of adult wildtype mice |  |  |  |  |  |  |  |  |  |
|  | SN |  |  |  |  | VTA |  |  |  |  |
| analyzed mRNA | mean | SD | SEM | median | N | mean | SD | SEM | median | N |
| β2a | 4.72 | 0.40 | 0.16 | 4.86 | 6 | 4.62 | 0.42 | 0.17 | 4.67 | 6 |
| β2e | 9.68*** | 1.76 | 0.72 | 10.25 | 6 | 8.43*** | 1.93 | 0.79 | 8.13 | 6 |

**Supplemental Table 5:**
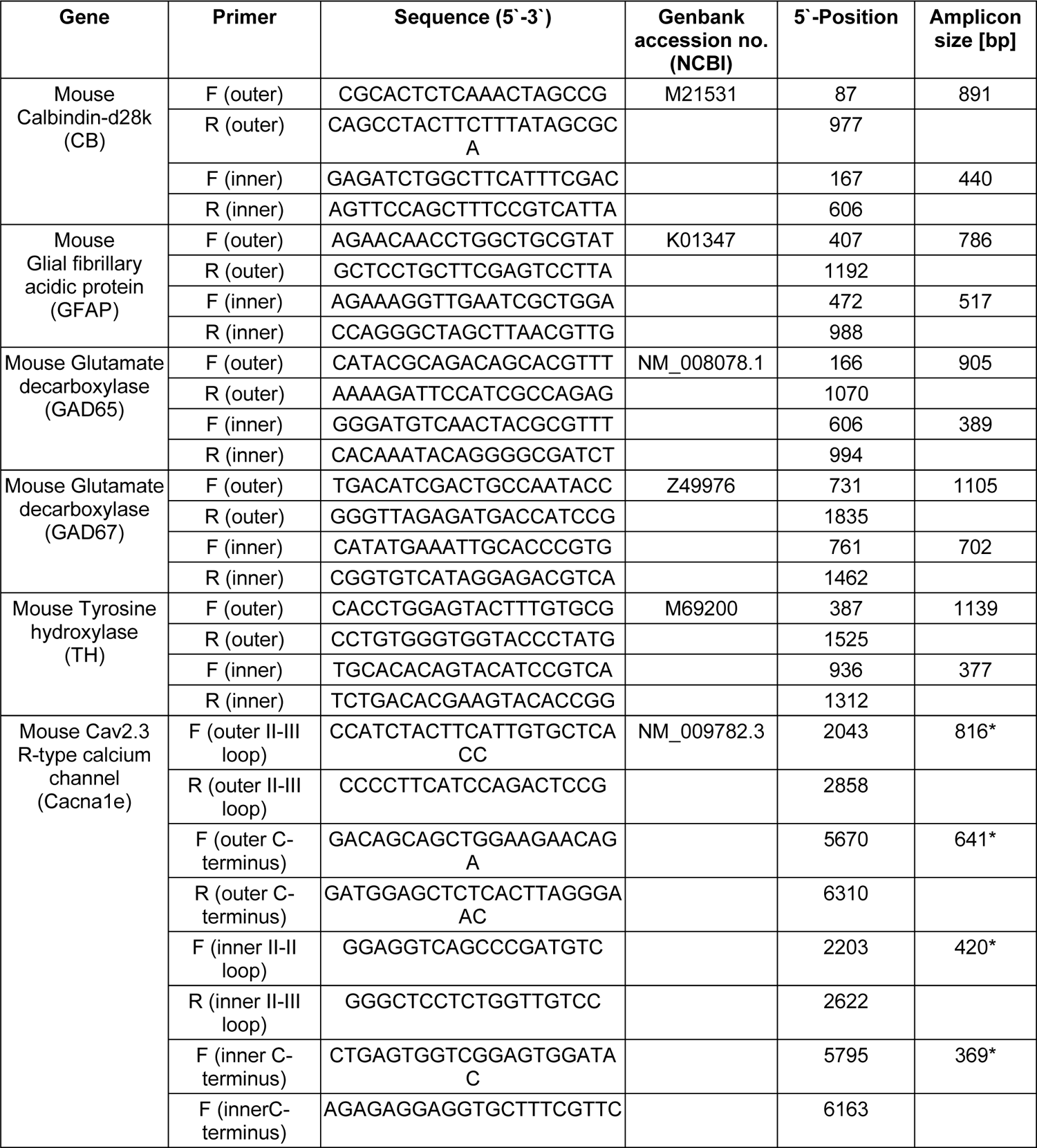
Multiplex PCR (outer) and nested PCR (inner) primers for qualitative PCR F: forward primer, R: reverse primer

**Supplemental Table 6.** Voltage-dependence of activation and inactivation of Cav1.3 co-transfected with α2δ1 and different β-subunits in tsA-201 cells All values are given as mean ± SEM for the indicated number of experiments (N=2). Parameters were obtained as described in Materials and Methods from a holding potential of −89 mV using 15 mM Ca^2+^ as the charge carrier. Voltage-dependence of gating: Parameters are as given in legend to Table 1. Statistical significance was determined using one-way ANOVA with Bonferroni post-hoc test (V_0.5_, V_rev_, act thresh, V_0.5,inact_, k_inact_, plateau) or Kruskal-Wallis followed by Dunn’s multiple comparison test (k). Statistical significances of post hoc tests are indicated for comparison vs. β2a (*, **, ***) or vs. β3 (^§^, ^§§^, ^§§§^): *** p<0.001; ** p<0.01; * p<0.05. Inactivation time course: The r values represent the fraction of I_Ca_ remaining after 50, 100, 250, 500, 1000 or 5000 ms during a 5-s pulse to V_max_. Statistical significance was determined using one-way ANOVA with Bonferroni post-hoc test. Statistical significances of post hoc tests are indicated for comparison vs. β2a: *** p<0.001; ** p<0.01; * p<0.05).

| β-subunit | Cav1.3<sub>L</sub> - Activation 15 mM Ca<sup>2+</sup> | | | | | | Cav1.3<sub>L</sub> - Inactivation 15 mM Ca<sup>2+</sup> | | | |
| --- | --- | --- | --- | --- | --- | --- | --- | --- | --- | --- |
|  | V<sub>0.5</sub> [mV] | k [mV] | V<sub>rev</sub> [mV] | act thresh [mV] | current density [pA/pF] | n | V<sub>0.5, inact</sub> [mV] | k<sub>inact</sub> [mV] | plateau [%] | n |
| β2a | 4.3<br>±1.8 | 10.0<br>±0.3 | 71.2<br>±2.0 | -34.5<br>±0.9 | -7.8<br>±0.6 | 14 | -18.3<br>±3.2 | 10.0<br>±1.0 | 36.2<br>±5.8 | 5 |
| c3S/c4Sβ2a | 0.2<br>±1.2 | 9.2*<br>±0.2 | 71.3<br>±1.1 | -34.9<sup>§§</sup><br>±0.9 | -8.4<br>±1.6 | 15 | -24.4<br>±1.1 | 6.1***<br>±0.4 | 21.6*<br>±2.1 | 11 |
| β3 | 3.1<br>±1.9 | 8.9*<br>±0.4 | 71.1<br>±2.7 | -30.6*<br>±0.8 | -10.2<br>±1.4 | 9 | -20.3<br>±0.9 | 7.0*<br>±0.5 | 28.5<br>±5.4 | 5 |
| Cav1.3<sub>L</sub> - 5 s Inactivation 15 mM Ca<sup>2+</sup> |  |  |  |  |  |  |  |  |  |  |
| β-subunit | r50 [%] | | r100 [%] | | r250 [%] | | r500 [%] | r1000 [%] | r5000 [%] | n |
| β2a | 86.8<br>±2.7 | | 81.2<br>±3.0 | | 71.5<br>±3.5 | | 63.3<br>±4.1 | 54.6<br>±4.2 | 36.0<br>±3.7 | 10 |
| c3S/c4Sβ2a | 74.5*<br>±3.0 | | 62.7***<br>±2.6 | | 47.1***<br>±3.2 | | 33.0***<br>±2.6 | 23.3***<br>±2.6 | 12.6***<br>±1.7 | 14 |
| β3 | 81.5<br>±3.4 | | 70.3<br>±3.7 | | 50.6**<br>±4.1 | | 34.6***<br>±3.9 | 25.2***<br>±4.2 | 13.7***<br>±3.0 | 7 |

**Supplemental Table 7:**
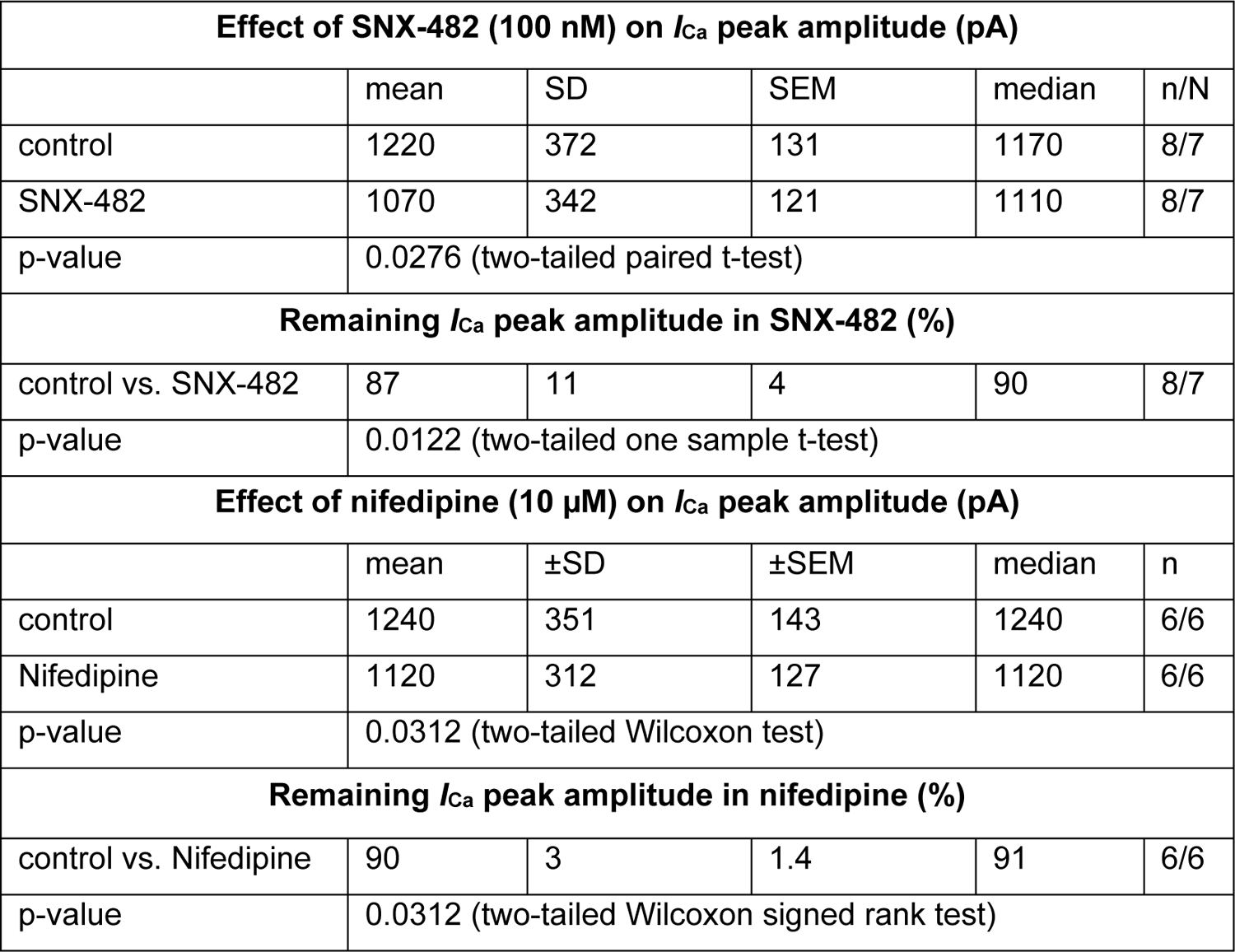
Inhibition of I_Ca_ in SN DA neurons of adult (12 weeks) mice by 100 nM SNX-482 and 10 µM nifedipine For details see results.

| Effect of SNX-482 (100 nM) on $I_{Ca}$ peak amplitude (pA) | | | | | |
| --- | --- | --- | --- | --- | --- |
|  | mean | SD | SEM | median | n/N |
| control | 1220 | 372 | 131 | 1170 | 8/7 |
| SNX-482 | 1070 | 342 | 121 | 1110 | 8/7 |
| p-value | 0.0276 (two-tailed paired t-test) |  |  |  |  |
| Remaining $I_{Ca}$ peak amplitude in SNX-482 (%) | | | | | |
| control vs. SNX-482 | 87 | 11 | 4 | 90 | 8/7 |
| p-value | 0.0122 (two-tailed one sample t-test) |  |  |  |  |
| Effect of nifedipine (10 μM) on $I_{Ca}$ peak amplitude (pA) | | | | | |
| | mean | ±SD | ±SEM | median | n |
| control | 1240 | 351 | 143 | 1240 | 6/6 |
| Nifedipine | 1120 | 312 | 127 | 1120 | 6/6 |
| p-value | 0.0312 (two-tailed Wilcoxon test) |  |  |  |  |
| Remaining $I_{Ca}$ peak amplitude in nifedipine (%) | | | | | |
| control vs. Nifedipine | 90 | 3 | 1.4 | 91 | 6/6 |
| p-value | 0.0312 (two-tailed Wilcoxon signed rank test) |  |  |  |  |

## Supplemental Methods

### Dissection of brain tissue for RT-qPCR

Tissue was dissected after mice had been sacrificed by cervical dislocation under isoflurane (Vetflurane, Vibac UK, 1000 mg/g) anesthesia. The freshly extracted mouse brains from 12-14 weeks old male mice were snap-frozen in isopentane (Carl Roth, catalog #3926.2) that was pre-cooled with dry ice (∼-40°C). To dissect VTA and SN tissue, 100 µm thick sections were cut on a cryostat (CM1950, Leica, Germany) and collected on glass coverslips. After cooling the coverslips on dry ice, regions of interest were punched under a dissection microscope using a pre-cooled sample corer (Fine Science Tools, Germany) (VTA: inner diameter 0.8 mm, 1 punch per hemisphere; SN: inner diameter 0.5 mm, 2 punches per hemisphere). For each brain region, tissue punches from both hemispheres of 7-8 successive 100 µm sections between Bregma −3.00 mm and −3.80 mm (according to Paxinos & Franklin, 2004) were collected in the sample corer. The tissue punches were transferred into an Eppendorf tube and again snap-frozen in liquid nitrogen. The punched brain sections were stained with cresyl violet (Nissl’s staining) for histological verification. Briefly, the slides with the punched sections were incubated in 4% PFA overnight. On the next day, the sections were rinsed in H_2_O and the following staining steps were performed: H_2_O 1 min, 70% ethanol 5 min, 100% EtOH 5 min, 0.5% cresyl violet solution pH 3.9 (2.5 g cresyl violet acetate, 30 ml 1M Na-acetate*3 H_2_O and 170 ml 1M acetic acid) 10 min, 70% ethanol 2 min, 100% ethanol 2 min, Xylol 1 min. The sections were coverslipped immediately with Eukitt (Sigma-Aldrich, catalog #03989), left to harden overnight and imaged on a bright field microscope at 2x.

### RNA isolation and cDNA synthesis for tissue RT-qPCR

RNeasy® Lipid Tissue Mini Kit (Qiagen GmbH, Germany, catalog #1023539) was used to isolate total RNA from brain tissues. Briefly, the tissue was disrupted and homogenized by vortexing samples for 5-10 minutes in 500 μl (SN, VTA) of phenol/guanidine-based Qiazol lysis reagent (Qiagen GmbH, Germany, catalog #79306) and passing the lysate ten times through a 21-gauge needle. All further steps, including purification of RNA with QIAGEN silica gel membrane technology (Qiagen GmbH, Germany), were performed according to the manufacturer’s protocol. For the final elution 15 μl of RNase-free water were used twice. An optional on-column DNase digestion (Qiagen GmbH, Germany, catalog #79254) was performed to reduce genomic DNA contamination. The RNA concentration was determined photometrically yielding approximately 20 ng/µl for VTA and SN or 1-2 µg/µl for whole brain.

RNA was reverse transcribed using Maxima H Minus First Strand cDNA synthesis kit with random hexamer primers following the manufactureŕs instructions (Thermo Fisher Scientific, Waltham, MA, USA). 1 µg or 13 μl of total RNA were used as template for reverse transcription. 1 µl cDNA corresponds to 0.65 x the amount of RNA equivalent.

### Standard curve method-based RT-qPCR for quantification of β-subunit expression in SN and VTA

In order to generate DNA templates of known concentrations for RT-qPCR standard curves, the concentration of the digested fragments was determined using the Quant-IT PicoGreen dsDNA Assay Kit (Invitrogen, Carlsbad, CA, USA). Subsequently, standard curves were generated using a serial dilution ranging from 10^7^ to 10^1^ DNA molecules in water containing 1 *μ*g/ml of poly-dC DNA (Midland Certified Reagent Company Inc., Midland, TX, USA). RT-qPCRs of standard curves and samples were performed as described previously (N. J. Ortner et al., 2020; Schlick et al., 2010). Samples for RT-qPCR quantification (50 cycles) contained 5 ng of total RNA equivalent of cDNA, the respective TaqMan gene expression assay, and TaqMan Universal PCR Master Mix (Thermo Fisher Scientific). Specificity of all assays was confirmed using different DNA ratios of corresponding and mismatched DNA fragments for the β1-4 and for the β2a-2e assays, respectively. Importantly, all assays specifically recognized the corresponding fragment even in the presence of a 10-fold higher concentration of other splice variants (Suppl. Fig. 1). RT-qPCR was performed in duplicates from three independent RNA preparations from three biological replicates. Samples without template served as negative controls. No RT RNA controls could not be included since all obtained RNA was transcribed into cDNA due to low yields. The expression of seven different endogenous control genes (*Actb*, *B2m*, *Gapdh*, *Hprt1*, *Tbp*, *Tfrc* and *Sdha*; N. J. Ortner et al., 2020; Schlick et al., 2010) was routinely measured and used for data normalization as previously described (Genome BiolVandesompele et al., 2002) (Suppl. Fig. 1, Suppl. Table 2). Briefly, data were normalized to the most stably expressed endogenous control genes (*Gapdh* and *Tfrc*) determined by geNorm (Suppl. Fig. 1). Normalized molecule numbers were calculated for each assay based on their respective standard curve. Standard curve parameters are given in (Suppl. Table 3). qPCR analyses were performed using the 7500 Fast System (Applied Biosystems, Foster Systems, CA, USA).

### Target probes for RNAScope *in situ* hybridization in identified SN DA and VTA DA neurons and image acquisition

Target probes were either obtained from the library of validated probes provided by Advanced Cell Diagnostics (ACD) or self-designed in cooperation with ACD to specifically discriminate between two splice variants of β2, namely β2a and β2e. Target probes (RNAScope assays) used for analysis were as follows:

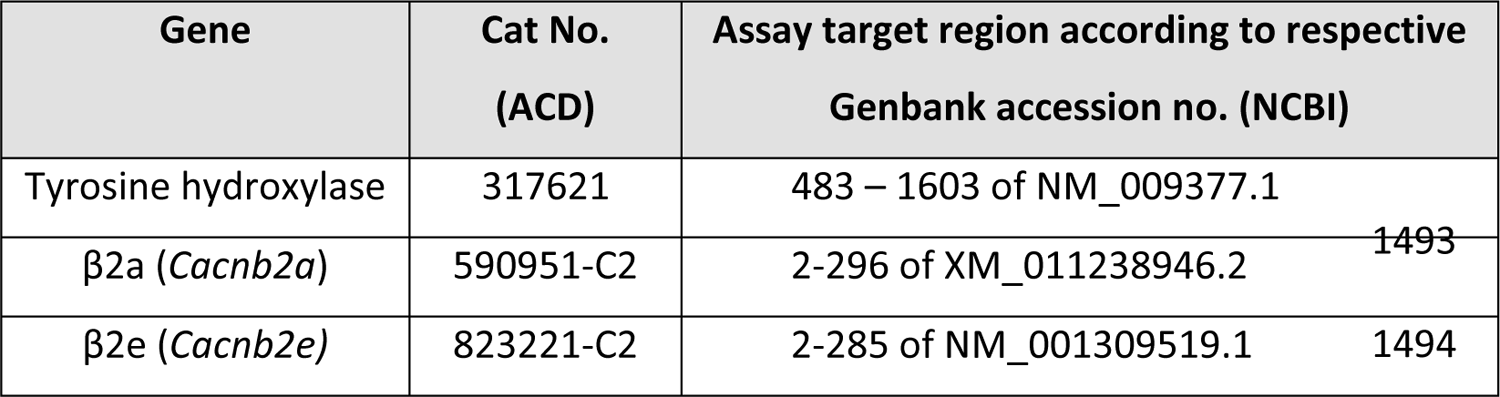

Target genes, visualized with Atto550 fluorophore, were co-stained with Tyrosine hydroxylase (TH), visualized with AlexaFluor488, as a marker for dopaminergic neurons. The gene peptidyl-prolyl isomerase B (PPIB) was used as positive control.

Fluorescent images of midbrain sections were acquired by a Leica CTR6 LED microscope using a Leica DFC365FX camera as z-stacks, covering the full depth of cells at 63x magnification. Z-stacks were reduced to maximum intensity Z-projections using Fiji (http://imagej.net/Fiji) and images were analyzed by utilizing a custom-designed algorithm (Wolution, Munich, Germany). The algorithm delineates cell shapes according to the TH marker gene signal and quantifies the area of fluorescent staining. According to Advanced Cell Diagnostics (ACD), target probe hybridization results in a small fluorescent dot for each mRNA molecule, allowing quantification of absolute number of mRNA molecules independent from fluorescent signal intensity.

### Multiplex-nested PCR, qualitative and quantitative PCR analysis in identified DA neurons

Cryosectioning, laser-microdissection, reverse transcription: Cryosectioning, UV-laser microdissection (UV-LMD) and reverse transcription were carried out similarly as previously described in detail (Benkert et al., 2019; Duda et al., 2018; Grundemann et al., 2011; Liss, 2002). Briefly, coronal 12 µm mouse midbrain cryosections were cut with a cryomicrotome CM3050 S (Leica), mounted on PEN-membrane slides (Mirodissect), stained with a cresyl-violet (CV) ethanol solution and fixed with an ascending ethanol series. UV-LMD of SN dopaminergic neurons from cresyl-violet-stained midbrain sections from juvenile mice was carried out using an LMD7000 system (Leica Microsystems). Reverse transcription was carried out directly without a separate RNA isolation step in a one-tube procedure.

All cDNA samples were precipitated as described (Liss, 2002). Qualitative multiplex-nested PCR and quantitative qPCRs were carried out essentially as described (Benkert et al., 2019; Duda et al., 2018; Grundemann et al., 2011). Briefly, qualitative multiplex-nested PCR was performed with the GeneAmp PCR System 9700 (Thermo Fisher Scientific) using 1/3 of each individual cDNA sample, corresponding to ∼3 SN DA neurons, for marker gene expression analysis: tyrosine hydroxylase (TH) as a marker for dopaminergic midbrain neurons, the glutamic acid decarboxylase isoforms GAD_65_ and GAD_67_ as markers for GABAergic neurons, the glial fibrillary acidic protein (GFAP) as a marker for astroglia cells and calbindin d28k (CBd28k), that is strongly expressed only in less vulnerable dopaminergic midbrain neurons. Only cDNA pools expressing the correct marker gene profile (i.e., TH positive, GAD, GFAP, CBd28k negative) were further analyzed via qualitative and quantitative PCR. Qualitative PCR for β2 splice variant expression analysis was performed with the QuantStudio 3 System (Thermo Fisher Scientific) using ∼3 SN DA neurons. Qualitative PCR products were analyzed in a QIAxcel Advanced System (Qiagen).

Details of multiplex PCR (outer) and nested PCR (inner) primers for qualitative PCR are shown in Suppl. Table 5.

For Cav2.3 splice variant analysis via qualitative RT-PCR, two cDNA fragments (mCav2.3 II-III loop and mCav2.3 C-terminus) covering the three respective splice sites were amplified in a duplex PCR, followed by two individual nested PCR reactions. PCR conditions: 15 min 95°C; 35 cycles: 30 s 94°C; 1 min 61°C; 3 min 72°C; 7 min 72°C for duplex PCR; and 3 min 94°C; 35 cycles: 30 s 94°C; 1 min 61°C; 1 min 72°C; 7 min 72°C for nested PCRs. Primer sequences are given in Supplementary Table 2. Outer duplex primer pairs were chosen using Oligo7.60 software (possible PCR amplicon sizes for mCav2.3 II-III loop: 816 bp, 795 bp and 759 bp; for mCav2.3 C-terminus: 770 bp and 641 bp). For the nested PCRs, primer sequences from Weiergraber et al., 2005 were used (possible PCR amplicon sizes for mCav2.3 II-III 980 loop: 420 bp, 399 bp and 363 bp; for mCav2.3 C-terminus: 498 bp and 369 bp). Nested PCR products were separated in a 4% agarose gel (MetaPhor agarose, Biozym), and expressed splice variants in individual SN DA neurons were identified according to the amplicon sizes of the nested PCR products (cDNA from mouse whole-brain tissue was used as positive control) as follows (II-III-loop + C-terminus): Cav2.3a: 363 + 369 bp; Cav2.3b: 399+369 bp; Cav2.3c: 420+369 bp; Cav2.3d. 420+498; Cav2.3e: 363 + 498 bp; Cav2.3f: 399 + 498 bp.

Quantitative real time PCR (qPCR) was performed with the QuantStudio 3 System (Thermo Fisher Scientific). qPCR assays (TaqMan®), were marked with a 3’ BHQ (black hole quencher) and 5’ FAM (Carboxyfluorescein). TaqMan assays were carefully established and performance was evaluated by generating standard curves, using defined amounts of cDNA (derived from midbrain tissue mRNA), over four magnitudes of 10-fold dilutions as templates, in at least three independent experiments, as described (Duda et al., 2018; Liss, 2002).

Details of all qPCR assays (TaqMan®), and standard curve parameters used for analysis are provided in Supplementary Table 4.

Relative qPCR quantification data are given as cDNA amount [pg/cell] with respect to midbrain-tissue cDNA standard curves, calculated according to the following formula, and were normalized to the respective cell size by dividing respective expression values to the corresponding area of the individual microdissected neurons.

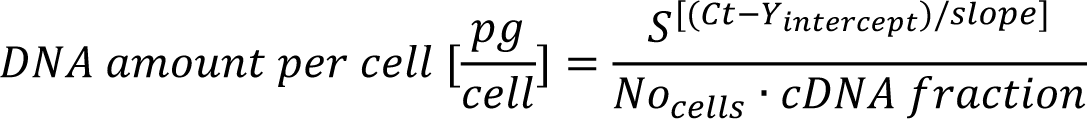

With S = serial dilution factor of the standard curve (i.e. 10), No_cells_ = number of SN DA neurons per sample (i.e. 10 here), cDNA fraction = fraction of the cDNA reaction used as template in the qPCR reaction (i.e. 5/17) and the Y-intercept and slope of the relative standards (see Supplementary Table 4). β2 TaqMan qPCR assay and standard curve information, used for relative qPCR-based transcript quantification: Note that the amplicon is very small (70 bp). The probe is 5’-FAM (6-carboxyfluorescein) and 3’-NFQ (non-fluorescent quencher) labelled. The assay Mm01333550_m1 (assay-ID) was used (assay location 583) spanning exon-boundary 4-5 of the mouse β2 gene (*Cacnb2*). Assay parameters were as follows: threshold for analysis: 0.05, Y-intercept of standard curve: 43.5 ± 0.2, slope: −3.43 ± 0.04, R^2^: 1.0 ± 0.0: means ± SEM, n=5.

### Slice preparation for perforated patch recordings in SN DA neurons

Animals were anesthetized with isoflurane (B506; AbbVie Deutschland GmbH and Co KG, Ludwigshafen, Germany) and subsequently decapitated. The brain was rapidly removed and a block of tissue containing the mesencephalon was immediately dissected. Coronal slices (250 − 300 μm) containing the SN were cut with a vibration microtome (HM-650 V; Thermo Scientific, Walldorf, Germany) under cold (4°C), carbogenated (95% O_2_ and 5% CO_2_), glycerol-based modified artificial cerebrospinal fluid (GACSF; Ye et al., 2006) containing (in mM): 250 glycerol, 2.5 KCl, 2 MgCl_2_, 2 CaCl_2_, 1.2 NaH_2_PO_4_, 10 HEPES, 21 NaHCO_3_, 5 glucose adjusted to pH 7.2 (with NaOH) resulting in an osmolarity of ∼310 mOsm. Brain slices were transferred into carbogenated artificial cerebrospinal fluid (ACSF). First, they were kept for 20 min in a 35°C ‘recovery bath’ and then stored at room temperature (24°C) for at least 30 min prior to recording. ACSF contained (in mM): 125 NaCl, 2.5 KCl, 2 MgCl_2_, 2 CaCl_2_, 1.2 NaH_2_PO_4_, 21 NaHCO_3_, 10 HEPES, and 5 Glucose adjusted to pH 7.2 (with NaOH) resulting in an osmolarity of ∼310 mOsm. SN dopaminergic neurons were identified according to their sag component / slow Ih-current, broad action potentials or post hoc by TH/DAT-immunohistochemistry (Lacey et al., 1989; Neuhoff et al., 2002; Richards et al., 1997). Biocytin-streptavidin labeling was combined with TH-immuno-histochemistry (Hess et al., 2013).

